# Feature selection and causal analysis for microbiome studies in the presence of confounding using standardization

**DOI:** 10.1101/2020.08.09.243188

**Authors:** Emily Goren, Chong Wang, Zhulin He, Amy M Sheflin, Dawn Chiniquy, Jessica E Prenni, Susannah Tringe, Daniel P Schachtman, Peng Liu

## Abstract

**Background:** Microbiome studies have uncovered associations between microbes and human, animal, and plant health outcomes. This has led to an interest in developing microbial interventions for treatment of disease and optimization of crop yields which requires identification of microbiome features that impact the outcome in the population of interest. That task is challenging because of the high dimensionality of microbiome data and the confounding that results from the complex and dynamic interactions among host, environment, and microbiome. In the presence of such confounding, variable selection and estimation procedures may have unsatisfactory performance in identifying microbial features with an effect on the outcome.

**Results:** In this manuscript, we aim to estimate population-level effects of individual microbiome features while controlling for confounding by a categorical variable. Due to the high dimensionality and confounding-induced correlation between features, we propose feature screening, selection, and estimation conditional on each stratum of the confounder followed by a standardization approach to estimation of population-level effects of individual features.

Comprehensive simulation studies demonstrate the advantages of our approach in recovering relevant features. Utilizing a potential-outcomes framework, we outline assumptions required to ascribe causal, rather than associational, interpretations to the identified microbiome effects. We conducted an agricultural study of the rhizosphere microbiome of sorghum in which nitrogen fertilizer application is a confounding variable. In this study, the proposed approach identified microbial taxa that are consistent with biological understanding of potential plant-microbe interactions.

**Conclusions:** Standardization enables more accurate identification of individual microbiome features with an effect on the outcome of interest compared to other variable selection and estimation procedures when there is confounding by a categorical variable.

## 1 Introduction

Advancements in next-generation sequencing (NGS) technologies have recently allowed for unprecedented examination of the community of microorganisms in a host or site of interest, referred to as a microbiome (Lederberg and Mccray, 2001). Early cultivation-dependent methods only allowed for detection of a small fraction of the total microbial species present. In contrast, NGS technologies can rapidly detect thousands of microbes in each sample by determining the nucleotide sequences of short microbial DNA fragments. These fragments may either correspond to targets of a specific genetic marker, commonly the 16S ribosomal RNA gene for taxonomic identification of bacteria as in amplicon sequencing, or result from shearing all the DNA in a sample as in shotgun metagenome sequencing (Riesenfeld et al., 2004). For each fragment, the corresponding nucleotide sequence is referred to as a “read,” the length of which is dependent on the specific NGS system (Liu et al., 2012).

Both amplicon-based and shotgun metagenomic approaches can enumerate the relative abundance of thousands of microbial features per sample. Use of amplicon sequencing for microbial enumeration is more common than shotgun metagenome sequencing due to reduced cost and complexity. For this reason, we focus on amplicon-based microbiome data here, and refer the reader to Sharpton (2014) for detailed coverage of metagenomic sequencing and Knight et al. (2018) for a thorough comparison of the two approaches. In order to enumerate microbes, amplicon reads are typically clustered into operational taxonomic units (OTUs) according to a fixed level of sequence similarity (e.g., 97%) (Westcott and Schloss, 2015), or as advocated by Callahan et al. (2017), enumerated on the basis of denoised sequences termed exact amplicon sequence variants (ASVs). Both OTUs and ASVs may be classified into known taxa (Schloss and Westcott, 2011). The resulting microbiome data for each sample are high-dimensional nonnegative integer counts across potentially thousands of features (taxa, OTUs, or ASVs). These counts represent relative, not absolute, numbers for each sample due to varying library sizes, a technical limitation of NGS approaches. Consequently, microbiome data must be normalized, rarefied, or treated as compositional in order to make comparisons across samples and it is unresolved which method is optimal for a particular research question and data set (Gloor et al., 2017; McMurdie and Holmes, 2014; Weiss et al., 2017).

Microbiome studies have uncovered associations between microbes and human, animal, and plant health outcomes. Randomized clinical trials have been performed to determine the causal effect of fecal microbiota transplantation (Camacho-Ortiz et al., 2017), but these do not provide causal inference on the contribution of individual microbiome features. It is important to identify individual microbiome features with a causal effect on the outcome because such discoveries may lead to development of microbial interventions for treatment of disease or optimization of crop yields. A recent review highlights the importance of identifying individual taxa with biologically relevant roles in microbiome studies (Banerjee et al., 2018).

Recently, there has been interest in causal inference in microbiome studies (Xia and Sun, 2017). The gold standard for causal inference is to randomly assign treatments (here, microbiome interventions) and estimate the causal effect. However, this is challenging in microbiome studies since many microorganisms cannot be directly cultured (Stewart, 2012), and random assignment of microbiomes to units is often not possible. To date, causal inference in microbiome studies has been primarily limited to causal mediation analysis that determines if a causal effect of treatment is transmitted through the microbiome (Sohn et al., 2019; Wang et al., 2019; Zhang et al., 2018). Software has been developed to apply Granger causality (Granger, 1969) to microbiome time series (Baksi et al., 2018), but the performance of such an approach has not been thoroughly evaluated using simulation studies.

In this work, we aim to identify individual microbial features with a causal effect on an outcome in a population of interest using causal inference. Here, the microbiome features are considered to be multivariate exposures, and are often of much higher dimension than the sample size. Previous work on high-dimensional causal inference is typically limited to settings with high-dimensional confounders rather than exposures (e.g., Schneeweiss et al. (2009)) or directed graphical modeling (Pearl, 2009). Recently, Nandy et al. (2017) considered directed graphical modeling for estimation of joint simultaneous interventions. However, their approach requires linearity and Gaussianity assumptions for high-dimensional inference, which are inappropriate for microbiome count data. There are proposed approaches for causal inference for multivariate exposures or treatments using the potential-outcomes framework, and such approaches often rely on the generalized propensity score (Imai and Van Dyk, 2004). Siddique et al. (2018) compared inverse probability of treatment weighting, propensity score adjustment, and targeted maximum likelihood approaches for multivariate exposures. Wilson et al. (2018) proposed Bayesian model averaging over different sets of confounders when the set of true confounding variables is unknown. When the exposures are time-varying, Taubman et al. (2009) considered *g*-estimation and Hernán et al. (2001) proposed a marginal structural model. However, in all of these studies with multivariate exposures, the exposure dimensionality is smaller than the sample size.

In addition to the high dimensionality, causal inference for microbiome studies is complicated by potentially complex interactions among host, environment, and microbiome. For example, there could be categorical confounding variables that affect both the outcome and some of the microbiome features. To overcome the challenges of the high dimensionality and presence of categorical confounding variables in microbiome studies, we propose standardization on the confounder and use the potential-outcomes framework for causal inference (Keiding and Clayton, 2014). The potential-outcomes framework (Holland, 1988; Neyman, 1923; Rubin, 1974) conceptually frames causal inference as a missing data problem: the outcome can only be measured under the exposure actually received, making the outcome unobservable under all other possible values of the exposure. We refer the reader to Hernán and Robins (2019) for a more detailed introduction. To deal with high-dimensionality of the microbiome exposure and categorical confounding variables, we propose variable screening, selection, and estimation of microbiome effects conditional on the confounder (i.e., stratification), followed by standardization to obtain estimates of effects in the population of interest. Conditioning on the confounder for microbiome feature screening, selection, and estimation avoids complications due to high marginal confounder-induced correlation between features. Further, conditional estimation naturally allows for effect modification (i.e., interaction between the confounder and microbiome features), affording flexibility to capture host-environment-microbiome interactions. Standardization allows for estimation and ranking of microbiome feature effects in the target population, which has policy and epidemiological relevance. Even if conditions for causal inference do not hold, avoiding such marginal correlation allows for superior identification of associational microbiome effects.

In this manuscript we begin by defining the estimands of interest and outlining conditions required for causal inference in Section 2. We then propose our estimation approach with standardization in Section 3. Next, we demonstrate the feasibility of our approach through simulation studies in Section 4 and present a real data application using an agricultural microbiome study in Section 5. This paper ends with a discussion and conclusion.

## 2 Model and assumptions

### 2.1 Notation, microbiome effects, confounding

Consider a study with *n* units (indexed by *i* = 1,…, *n*) aimed at identifying the population effect ***β*** = (*β*_1_,…, *β_p_*)′ of *p* microbiome features (e.g., taxa, ASVs, OTUs) ***A**_i_* = (*A*_*i*1_,…, *A_ip_*)′ on an outcome *Y_i_* ∈ ℝ. For formulating the estimand, we assume that ***A**_i_* has been appropriately normalized. Importantly, *Y_i_* represents the observed outcome, which differs from the notion of a potential outcome (Rubin, 1974). Define the potential outcome 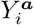 as the value the outcome would take under the (possibly counterfactual) microbiome value ***a*** = (*a*_1_,…, *a_p_*)′. Assume that the expected potential outcome is related to the population effect ***β*** through a linear function of the microbiome features as

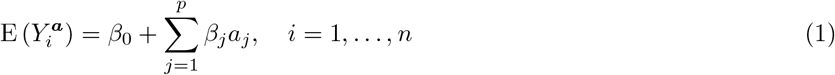

where for each *j*, *β_j_* represents the effect of the *j*th microbiome feature in the population. In terms of (1), identifying which microbiome features have a causal effect on the response corresponds to estimation and inference for *β_j_* (1 ≤ *j* ≤ *p*). For generality, the formulation of Equation (1) ignores possible microbe-microbe interactions and any constraints of carrying capacity.

Note that the model in (1) is defined for the potential outcomes, not the observed data, and is thus a marginal structural model (Hernán et al., 2001). In the presence of a confounding variable *L_i_* that affects both ***A**_i_* and *Y_i_*, this model generally does not hold for the observed data because confounding implies 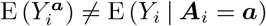. Consequently, specific assumptions and methodology are required to obtain an estimator 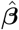 of ***β*** that has causal, rather than merely associational, interpretation. In the next sub-section, we address the assumptions required for such a causal interpretation. We restrict our attention to the case where the confounder *L_i_* is categorical with a finite number of levels, each represented sufficiently in the study of *n* units.

### 2.2 Assumptions for causal inference

Under the potential-outcomes framework, ascribing a causal interpretation to an estimate of ***β*** requires three assumptions: positivity, conditional exchangeability, and consistency (Hernán and Robins, 2019). Positivity requires positive probability for each possible microbiome level, conditional on the confounder. To formalize this, let 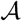 denote the set of all possible microbiome values in the population. The positivity condition holds if Pr (***A**_i_* = ***A*** | *L_i_* = *l*) > 0 for all 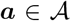 and all levels *l* of confounder *L_i_* such that Pr(*L_i_* = *l*) ≠ 0 in the population of interest, henceforth denoted by the set 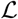. Clearly, if a given microbe is either absent or below the limit of detection across all samples, its effect on the response cannot be determined. Hence, this assumption requires a large enough sequencing depth in order to sufficiently enumerate any present microbes with a causal effect. Practical considerations for evaluating the positivity assumption are covered by Westreich and Cole (2010).

To meet the conditional exchangeability requirement, the data-generating mechanism for each possible microbiome must depend only on the confounder, formalized as 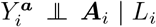 for all 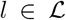, where ╨ denotes statistical independence. Conditional exchangeability requires no unmeasured confounding. This assumption is most justifiable in experiments where the confounder is randomly assigned as in our motivating study described later in Section 5, where agricultural plots are randomized to either low or high nitrogen fertilizer.

The consistency criterion is met if the observed outcome for each unit is the potential outcome under the observed microbiome, formally stated as 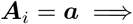 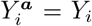. For microbiome data, this necessitates appropriate normalization. Since NGS-based technologies enumerate based on genetic material, the resulting counts can arise from both viable and non-viable microbes (Boers et al., 2016). In order to met the consistency assumption, relevant microbes with the same normalized count cannot have disparate effects due to differential viability. When there is concern that this assumption may be violated, it is possible to restrict amplification of RNA target genes to only viable bacterial cells (Rogers et al., 2008). We note that even if these three conditions cannot be verified, our proposed method has utility in estimation of associational, rather than causal, effects.

## 3 Methods

### 3.1 Standardization

Our goal is to estimate the population microbiome effects ***β*** of (1) and infer which microbiome features are relevant to the response, that is, {1 ≤ *j* ≤ *p*: *β_j_* ≠ 0}. We propose computing an estimate 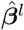 for each stratum 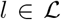 of the confounder, followed by standardization to the confounder distribution, thereby obtaining a population-level estimate 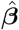. Under the assumptions stated in Section 2.2, there is no confounding within each stratum *l* of the confounder. Beyond elimination of confounding, conditioning on a stratum of the confounder avoids marginal correlation between features induced by the relationship with the confounder that can hinder feature selection performance. Figure S7 in the Supplementary Materials shows microbiome data from an agricultural study described in Section 5 where many features are highly correlated when considered marginally, but are relatively uncorrelated within each level of a fertilizer confounder. Combining the assumptions of Section 2.2 with the model in (1) and allowing for effect modification, we have

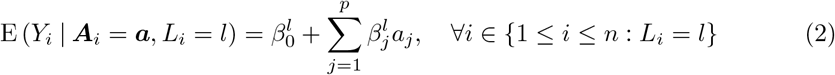

where 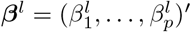 is the corresponding stratum-specific effect. There is effect modification if ***β**^l^* ≠ ***β**^l′^* for some 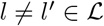.

Standardizing the stratum-specific mean outcomes to the confounder distribution produces the population mean outcome function

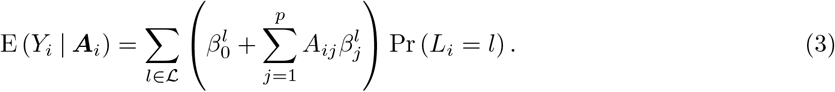

By linearity, the effect in the population corresponding to a one-unit increase in the *j*th microbiome feature, controlling for all others, is represented by 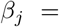 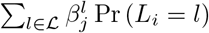 for *j* = 1,…, *p*. Given a suitable estimator 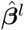 of ***β**^l^* for all 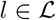, the resulting population-standardized estimate of *β_j_* is

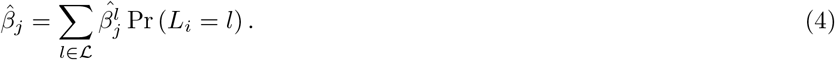

### 3.2 Feature selection and estimation

In this section, we propose a feature selection and estimation procedure for stratum-specific coefficients ***β**^l^*, performed independently for each confounder level 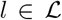. Within each stratum, we make a sparsity assumption that few microbiome features have an effect on the response and correspondingly most entries of ***β**^l^* are zero, and also assume that the outcome is normally distributed with constant variance. Commonly, *n* ≪ *p* for microbiome features for taxa at the level of species (and perhaps genera), OTUs, or ASVs. Consequently, we suggest penalized least squares estimation that induces shrinkage towards zero via a penalty function *p_λ_*, where *λ* is a tuning parameter controlling the amount of shrinkage. We suggest choosing *λ* using the Bayesian information criterion (BIC) (Schwarz, 1978) due to its consistency property in selecting the true features in certain settings (Wang et al., 2007) and nonconsistency of prediction accuracy criteria such as cross-validation (Leng et al., 2006). Possible choices for penalties that perform variable selection through shrinkage-induced sparsity include the least absolute shrinkage and selection operator (LASSO) (Tibshirani, 1996) and smoothly clipped absolute deviation (SCAD) (Fan and Li, 2001), among others (Zhang et al., 2010).

Due to the high dimensionality of microbiome data, variable screening in conjunction with penalized estimation may improve accuracy and algorithmic stability (Fan and Lv, 2008). The sure independence screening (SIS) of Fan and Lv (2008) retains features attaining the highest marginal correlation with the response, which may lead to poor performance when irrelevant features are more highly correlated with the response, marginally, than relevant ones. Since this is likely the case for microbiome data, we instead consider using the iterative sure independence screening procedure proposed by Fan and Lv (2008) and implemented by Saldana and Feng (2018) that avoids such a drawback by performing iterative feature recruitment and deletion based on a given penalty *p_λ_*.

### 3.3 Post-selection inference and error rate control

Inference on which microbiome features have a population-level effect, conducted by testing the null hypothesis *H*_0*j*_: *β_j_* = 0 for the *j*th feature (1 ≤ *j* ≤ *p*), is challenging using penalized least squares estimation. For example, the asymptotic distribution of the LASSO may not be continuous and is difficult to characterize in high-dimensional settings (Knight and Fu, 2000). Many approaches for error rate control post-variable selection using penalized regression make use of data splitting techniques (Bühlmann et al., 2014; Dezeure et al., 2015) but have low power for the small sample sizes common to microbiome studies. Due to these reasons, for inference we propose using the debiased, also known as desparsified, LASSO (van de Geer et al., 2014; Zhang and Zhang, 2014) applied to the estimate 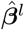 obtained using the LASSO penalty with the iterative SIS procedure. To make the computation tractable, we only apply the debiasing procedure to the features not screened out by the iterative SIS procedure and let 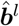 denote the resulting estimate. Under regularity assumptions and appropriate penalization, the debiased LASSO estimator has a limiting normal distribution (Dezeure et al., 2015).

For the *j*th feature, the standardized debiased iterative SIS-LASSO estimate 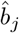 and its standard error are given by

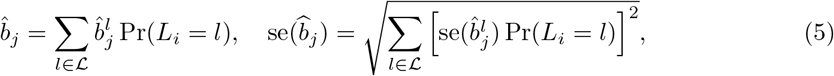

respectively, where the standard error formula follows from the independence of the strata. To obtain an estimator of the standard error, we plug-in the estimate 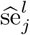 of 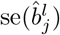 given by Dezeure et al. (2017) under homoscedastic errors if the *j*th feature was not removed by screening in the *l*th confounder stratum. We compute a *p*-value for testing *H*_0*j*_: *β_j_* = 0 versus *H*_1*j*_: *β_j_* ≠ 0 according to 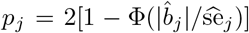 if feature *j* was not screened out in all confounder strata for *j* = 1,…, *p*, where Φ(·) denotes the standard normal cdf. To control the false discovery rate (FDR), we apply the Benjamini-Hochberg (BH) adjustment across all *p* features (Benjamini and Hochberg, 1995) to account for multiplicity in all features, including those that were removed from all strata.

## 4 Simulation Studies

Here, we evaluate our proposed standardization method using simulation studies. The simulation settings were designed to mimic microbiome studies seen in practice. To emulate species-level data, we consider *p* = 2, 000 microbiome features. To reflect data summarized at the genus level, we also consider *p* = 50. We consider sample sizes of *n* = 50 and *n* = 100, and assume the confounder is a binary indicator that takes the value one for *i* = 1,…, *n*/2 and zero for *i* = *n*/2 + 1,…, *n*.

### 4.1 Data-generating model for microbiome features

Conditional on the confounder *L_i_* = 0, the count data for the *j*th microbiome feature were drawn independently from a negative binomial distribution with mean *γ*_0*j*_ and dispersion *φ_j_* parameterized such that Var(*A_ij_*) = *γ*_0*j*_ + *φ_j_*(*γ*_0*j*_)^2^. That is, when *L_i_* = 0, the baseline mean for feature *j* is *γ*_0*j*_. When the confounder is present (*L_i_* = 1), the microbiome feature counts were drawn independently from a negative binomial distribution with mean *γ*_0*j*_*γ*_1*j*_ and dispersion *φ_j_*. Hence, *γ*_1*j*_ represents the multiplicative change in the mean relative to when the confounder is absent. If *γ*_1*j*_ ≠ 1, then feature *j* is affected by the confounder and otherwise *γ*_1*j*_ = 1. The first 30% of features were set to be affected by the confounder (differentially abundant between condition *L_i_* = 0 and condition *L_i_* = 1). More specifically, we simulated parameters *γ*_0*j*_ and *γ*_1*j*_ from the following distributions for *j* = 1,…, *p*:

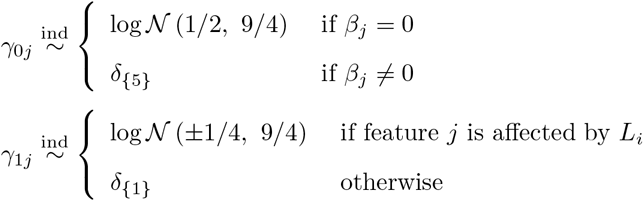

where *δ*_{*x*}_ represents a point mass at *x*. Our rationale for setting the baseline mean to five for relevant features (*β_j_* ≠ 0) was to ensure that they were sufficiently abundant for feature selection. We set the dispersions *φ_j_* = 10^−1^ for all features *j* = 1,…, *p* and simulated the microbiome count data ***A**_i_* with negative binomial distributions. In addition, we conducted a second set of simulations with *φ_j_* = 10^*−*6^, which approximates a Poisson distribution.

### 4.2 Data-generating model for response

Given the confounder and microbiome features ***A**_i_* simulated from the above sub-section, we draw the responses independently from a normal distribution with mean *μ_i_*(***Ã***_*i*_, *L_i_*) and variance *σ*^2^, where ***Ã***_*i*_ represents ***A**_i_* after centering and scaling (to mean zero and variance one within strata) and

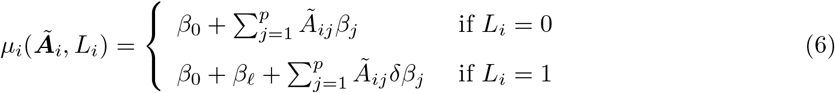

for *i* = 1,…, *n*. For more intuitive comparison of effect modification size, model (6) has an additive effect *β_1_* for the intercept and multiplicative effect *δ* for microbiome feature effects when *L_i_* = 1 compared with *L_i_* = 0. In particular, *β_1_* represents the direct confounder effect and *δ* is an effect modification parameter. Our simulation considers the case when there is no effect modification (*δ* = 1) as well as strong effect modification (*δ* = −0.9) where the relevant microbiome effects are large within each level of the confounder but small overall in the population. The response variability was set to *σ*^2^ = 1/16 for all scenarios. A total of *s* = 5 features were set to be relevant, with the non-zero elements of ***β*** set to (3, −3, 3, −3, 3). Our motivation for setting |*β_j_*| = 3 for all relevant *j* is to ensure the *β*_min_ property for model selection consistency is met within all strata for all simulation scenarios (Bühlmann et al., 2014). The choice of *s* = 5 yields sparsity such that *s* < *n_l_*/*log*(*p*) for most, but not all, simulation scenarios. Three scenarios covering differing proportions of the relevant features set to be confounded (*β_j_* ≠ 0 and *γ*_1*j*_ ≠ 1) were considered: either all (100% confounded), the first three (60% confounded), or none (0% confounded). To summarize our simulation settings, we have considered two dimensions of microbiome features: *p* = 2, 000 and *p* = 50; two sample sizes: *n* = 50 and *n* = 100; two distributions of microbiome count data: negative binomial and Poisson; inclusion of effect modifier: none or strong effect modifier; and three different proportions of confounded relevant features: 100%, 60%, and 0%. Hence, in total, we examined 48 different simulation settings. For each simulation setting, a total of 100 data sets were simulated.

### 4.3 Screening, penalization, and comparison models

We denote our proposed approach of estimation conditional on each stratum followed by standardization as “Conditional Std”. We investigate the performance of variable section using the LASSO and SCAD penalties for *p_λ_* both with and without screening, as well as the proposed inference procedure using the debiased LASSO with iterative SIS described in Section 3.3.

We compare our approach with six other models applied to the pooled data set, as opposed to conditionally on each stratum. The six comparison models are constructed based on three inclusion strategies for the confounder effect *β_1_* of Equation (6) and two possibilities for modeling effect modification. The confounder effect is either subject to screening and variable selection (“Select L”), forced to be included without penalization (“Require L”), or removed from the model entirely (“Ignore L”). We either model each microbiome feature effect as common across all confounder strata (corresponding to models with the aforementioned names) or allow for effect modification through stratum-specific microbiome feature effects denoted with the suffix “EffMod.” For each of the six models under comparison, we also investigate the performance of variable section using the LASSO and SCAD penalties for *p_λ_* both with and without screening, as well as the proposed inference procedure using the debiased LASSO with iterative SIS.

Table 1 presents the objective function for our proposed “Conditional Std” approach and the other six models under comparison. For the proposed approach “Conditional Std,” screening is based on iterative SIS recommended defaults applied to each stratum, whereas for all other approaches it is applied to the entire data set to correspond with the assumed model, resulting in different maximum model sizes shown in Table 2. The variables considered in the iterative SIS procedure for each model detailed in Table 2 correspond to those penalized in the objective function in Table 1. For “Conditional Std” and models allowing effect modification (suffix “EffMod”), the population estimates are computed according to Equation (4). These models center and scale each microbiome feature within each stratum, denoted by *Ã_ij_*. For models that do not allow for effect modification, the microbiome features are centered and scaled to have mean zero and variance one across all observations, regardless of stratum, denoted by 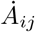.

**Table 1.** Models considered in simulation studies using penalized regression (with penalty *p_λ_*) for a binary confounder *L_i_* ∈ {0, 1}. *Ã_ij_* denotes microbiome feature *j* centered and scaled within each stratum; 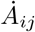 denotes microbiome feature *j* centered and scaled across all observations, regardless of stratum.

| Model | Objective function |
| --- | --- |
| Conditional Std | $\sum_l \frac{1}{2n_l} \sum_i I(L_i = l) \left( y_i - \beta_0^l - \sum_j \tilde{A}_{ij} \beta_j^l \right)^2 + \sum_{l,j} p_{\lambda_l}(\beta_j^l)$ |
| Select L | $\frac{1}{2n} \sum_i \left( y_i - \beta_0 - L_i \beta_\ell - \sum_j \dot{A}_{ij} \beta_j \right)^2 + \sum_j p_\lambda(\beta_j) + p_\lambda(\beta_\ell)$ |
| Select L EffMod | $\frac{1}{2n} \sum_i \left( y_i - \beta_0 - L_i \beta_\ell - \sum_{l,j} I(L_i = l) \tilde{A}_{ij} \beta_j^l \right)^2 + \sum_l \sum_j p_\lambda(\beta_j^l) + p_\lambda(\beta_\ell)$ |
| Require L | $\frac{1}{2n} \sum_i \left( y_i - \beta_0 - L_i \beta_\ell - \sum_j \dot{A}_{ij} \beta_j \right)^2 + \sum_j p_\lambda(\beta_j)$ |
| Require L EffMod | $\frac{1}{2n} \sum_i \left( y_i - \beta_0 - L_i \beta_\ell - \sum_{l,j} I(L_i = l) \tilde{A}_{ij} \beta_j^l \right)^2 + \sum_l \sum_j p_\lambda(\beta_j^l)$ |
| Ignore L | $\frac{1}{2n} \sum_i \left( y_i - \beta_0 - \sum_j \dot{A}_{ij} \beta_j \right)^2 + \sum_j p_\lambda(\beta_j)$ |
| Ignore L EffMod | $\frac{1}{2n} \sum_i \left( y_i - \beta_0 - \sum_{l,j} I(L_i = l) \tilde{A}_{ij} \beta_j^l \right)^2 + \sum_l \sum_j p_\lambda(\beta_j^l)$ |

**Table 2.** Variable screening and selection for models considered in simulation studies for a binary confounder *L_i_* ∈ {0, 1}.

| Model | Variables screened | Maximum model size |
| --- | --- | --- |
| Conditional Std | $\{\beta_1^l, \dots, \beta_p^l\}$ (independently $\forall l$) | $d_l = \lfloor n_l / \log(n_l) \rfloor \forall l$ |
| Select L | $\{\beta_1, \dots, \beta_p, L\}$ | $d = \lfloor n / \log(n) \rfloor$ |
| Select L EffMod | $\{\beta_1^{l=0}, \dots, \beta_p^{l=0}, \beta_1^{l=1}, \dots, \beta_p^{l=1}, L\}$ | $d = \lfloor n / \log(n) \rfloor$ |
| Require L | $\{\beta_1, \dots, \beta_p\}$ (given $L$) | $d = \lfloor n / \log(n) \rfloor$ |
| Require L EffMod | $\{\beta_1^{l=0}, \dots, \beta_p^{l=0}, \beta_1^{l=1}, \dots, \beta_p^{l=1}\}$ (given $L$) | $d = \lfloor n / \log(n) \rfloor$ |
| Ignore L | $\{\beta_1, \dots, \beta_p\}$ | $d = \lfloor n / \log(n) \rfloor$ |
| Ignore L EffMod | $\{\beta_1^{l=0}, \dots, \beta_p^{l=0}, \beta_1^{l=1}, \dots, \beta_p^{l=1}\}$ | $d = \lfloor n / \log(n) \rfloor$ |

### 4.4 Results

Simulation performance was summarized across all 100 simulated data sets for each scenario, model, and variable selection method considered using the true positive rate (TPR) and false positive rate (FPR). Given the selected variables, TPR measures the proportion of relevant features detected, while FPR measures the proportion of irrelevant features declared to be relevant, and these are computed here at the population-level by

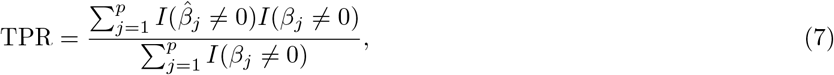

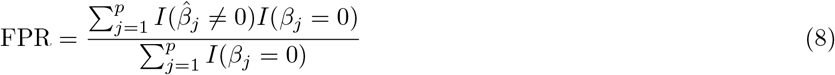

for all methods except the debiased LASSO inference procedure where 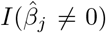 is replaced with the decision rule induced by the corresponding hypothesis test with FDR control at 0.05. An ideal method would take (TPR, FPR) values (1, 0). Supplementary Material Table S1 shows the average TPR and FPR across the 100 simulated data sets for the 12 simulation settings with Poisson distributed features and *n* = 100. The table lists the results for our proposed approach “Conditional Std” model and the other six models under comparison across different variable selection methods. Generally, the proposed “Conditional Std” model performed better than other models applied to the entire data set across different variable selection methods considered. When effect modification is present, the proposed approach has the highest mean TPR and lowest mean FPR for both the LASSO and SCAD penalties, both with and without screening, often achieving perfect rates on average. For the debiased LASSO applied after iterative SIS with the BH procedure and FDR control set to 0.05 (denoted by “iterSIS-dbLASSO-BH”), the proposed approach has the highest TPR and among the lowest FPR under strong effect modification across variable selection methods. This is not the case only when no effect modification is present, under high dimensionality (*p* = 2, 000), and not all relevant features are not confounded.

For post-selection inference based on the debiased LASSO following screening with iterative SIS, we evaluated the area under the receiver operating characteristic curve (AUC) using the *p*-values for testing *H*_0*j*_: *β_j_* = 0 as the classifiers. AUC aggregates classification performance of TPR versus FPR across different classification thresholds, taking the value 1 for perfect prediction, 0.5 for random guessing, and 0 for always wrong prediction. Box plots of the AUC across 100 data sets for each model are shown in Figure 1 for 12 simulation settings with *n* = 100 and Poisson features (results for *n* = 50 and negative binomial features are presented in Supplementary Material Figures S1–S3). The proposed approach has near perfect ranking under low dimensionality (*p* = 50) for all settings and under high dimensionality (*p* = 2, 000) when all relevant features are impacted by the confounder. Similar to the results in Supplementary Material Table S1, the proposed approach performs best out of all models considered except when effect modification is not present and at least some relevant features are not confounded.

**Figure 1.**
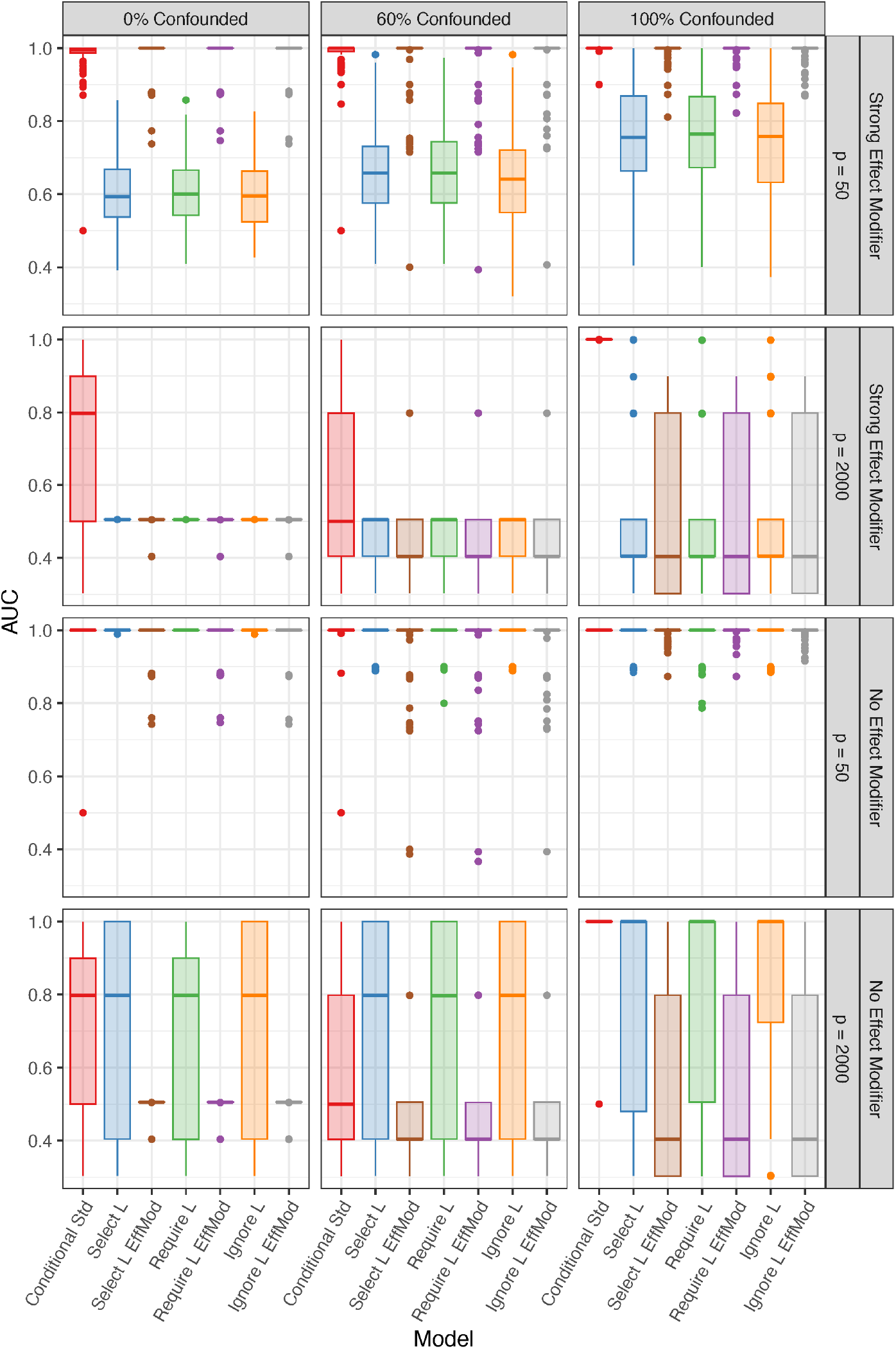
Simulation results. Box plots of the area under the curve (AUC) from 100 simulation replications for *n* = 100 and Poisson features using *p*-values based on the debiased LASSO estimate following iterative sure independence screening.

To evaluate false discovery rate (FDR) control for varying thresholds *α* = (0.01, 0.02,…, 0.10) commonly used in practice, we computed the false discovery proportion (FDP) at a given *α* value for debiased LASSO inference according to

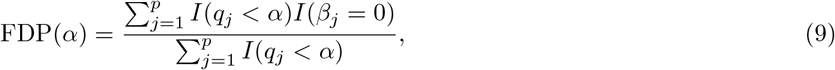

where *q_j_* is the BH-adjusted *p*-value (or *q*-value) for feature *j*. A well performing model will have FDP(*α*) ≤ *α*. For *n* = 100 and Poisson features, Figure 2 shows that the proposed “Conditional Std” model appropriately controls FDR under low dimensionality (*p* = 50). For high dimensionality (*p* = 2, 000), the proposed approach does not control FDR when at least some relevant features are not confounded, though the observed mean FDP does not exceed the nominal level greatly when compared to other competing models applied to the pooled data. The FDR control for the other six models under comparison is either very conservative or highly liberal. Similar results were seen for *n* = 50 and negative binomial features, though lack of FDR control was more common for the *n* = 50 case (Supplementary Materials Figures S4–S6).

**Figure 2.**
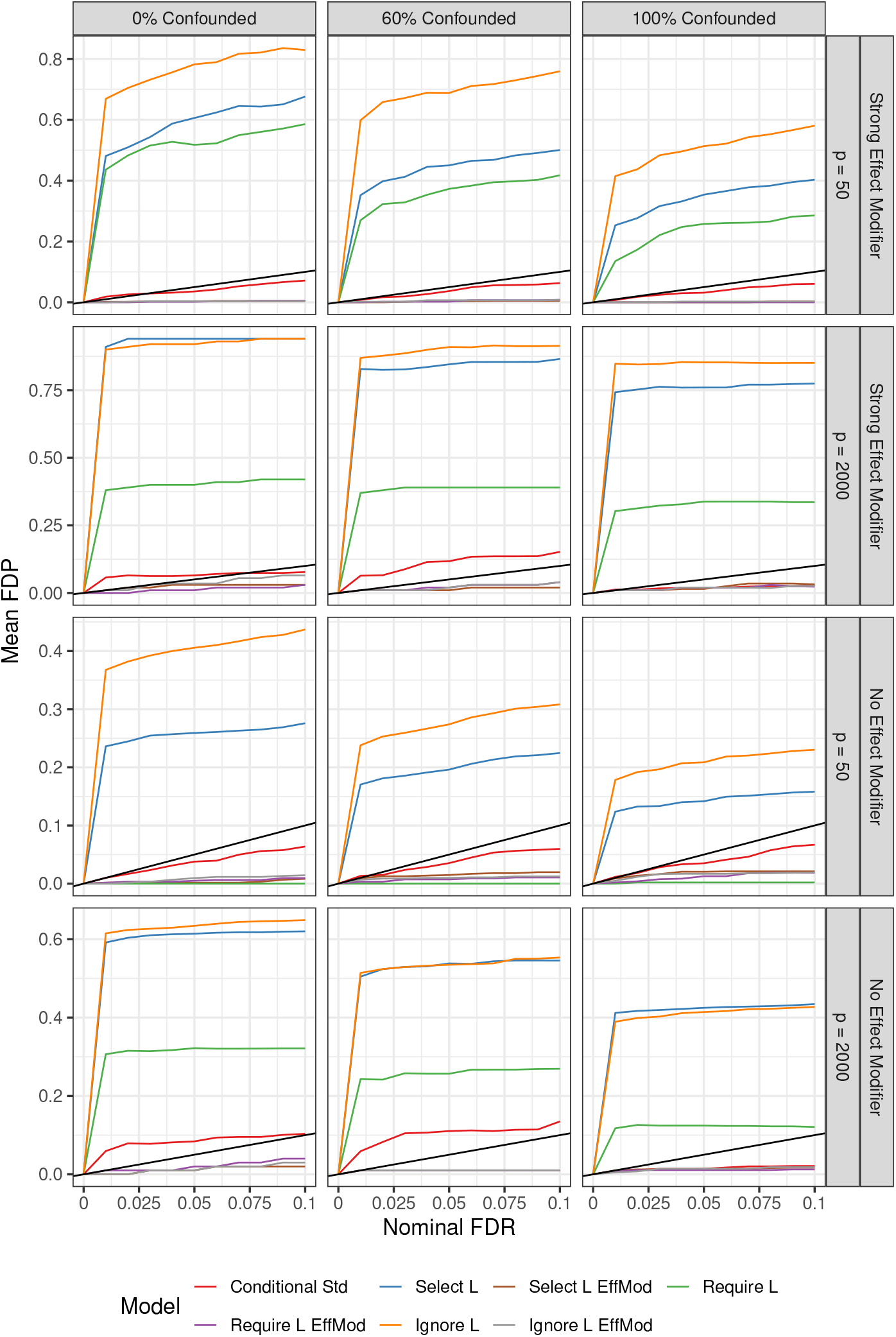
Simulation results. Mean estimated false discovery proportion (FDP) for *n* = 100 and Poisson features at varying nominal false discovery rate (FDR) values using Benjamini-Hotchberg adjusted *p*-values based on the debiased LASSO estimate following iterative sure independence screening (iterative SIS). The *y* = *x* line is shown in black; any values above this line indicate lack of FDR control.

## 5 Real Data Analysis

We conducted a microbiome study to investigate the effect of the rhizosphere microbiome of the cereal crop sorghum (*Sorghum bicolor*) on the phenotype 12-oxo phytodienoic acid (OPDA) production in the root. Sorghum root production of OPDA is of primary interest due to OPDA having both independent plant defense functions and being an important precursor to Jasmonic acid, which functions in plant immune responses that are induced by beneficial bacteria (Van der Ent et al., 2009; Wasternack, 2014). The study analyzed here is part of an experiment described by Sheflin et al. (2019); we subset on *n* = 34 samples collected in September across high and low nitrogen fertilizer. Rhizosphere microbiome data were collected using 16S amplicon sequencing and clustered at 97% sequence identity. The resulting 5, 584 OTUs were rarefied to 20, 000 reads per observation and low abundance OTUs (less than 4 non-zero observations out of 34) were excluded (Xiao et al., 2018), leaving a total of 4, 244 OTUs.

Pairwise Spearman’s correlations for the feature counts are shown in Figure S7 in the Supplementary Materials for the 150 largest marginal correlations (pooling samples over nitrogen fertilizer levels), which contrast to the small correlations within nitrogen stratum. Using our proposed procedure of testing the standardized feature effect using the debiased LASSO following iterative SIS applied to each nitrogen level, a total of four microbiome features with an effect on root ODPA production were identified while FDR was controlled at 0.05 with BH adjustment (Table 3). Nitrogen stratum-specific residuals did not indicate any violation of the assumptions of constant variance or normality (Figures S8–S9 of the Supplementary Materials).

**Table 3.**
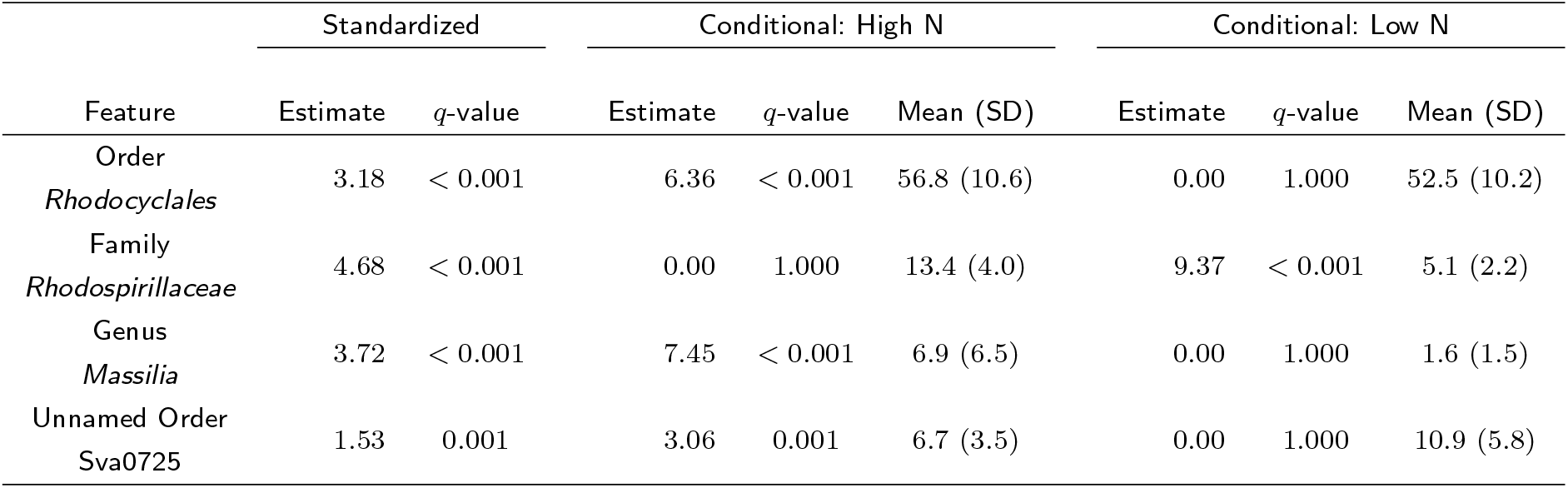
Sorghum study analysis results: features with a significant effect on sorghum root ODPA production in the study population with FDR control at the 0.05 level using the Benjamini-Hochberg (BH) procedure on the debiased LASSO estimate following sure independence screening (iterative SIS). The corresponding conditional estimates and rarefied mean abundance (standard deviation) is also presented.

Each microbiome feature effect identified at the study population-level was only identified in one nitrogen condition, though abundance did not differ greatly between the two nitrogen strata (Table 3). Specifically, only one feature was estimated to be more abundant under low nitrogen, and this feature was classified as belonging to the *Rhodospirillaceae* family (nonsulfur photosynthetic bacteria), of which nearly all members have the capacity to fix molecular nitrogen (Madigan et al., 1984). Various strains of *Rhodospirillaceae* have shown potential to promote plant growth in the grass species *Brachiaria brizantha* (Silva et al., 2013). Consequently, the increased levels of root OPDA content may have been the result of bacterial synthesis (Forchetti et al., 2007). While less is known about the three additional significant features, the overall findings are in alignment with biological understanding of potential plant-microbe interactions.

## 6 Discussion

We have proposed and evaluated methodology for causal inference for individual features in high-dimensional microbiome data using standardization. These techniques are typically employed in epidemiology and use the potential-outcomes framework, in contrast to graphical models, which are a more common approach for high-dimensional causal inference but usually require Gaussian assumptions for inference that are often violated by microbiome data (Pearl, 2009). Instead, our approach conditions on the confounder and shows favorable results for Poisson and negative binomial microbiome features. Compared to estimation methods applied to the entire data set, the proposed standardization approach typically demonstrated superior recovery of relevant microbiome effects accross multiple variable screening and selection procedures.

Association and causation are not equivalent even for a one-dimensional treatment or exposure, and the challenges of causal analysis are exacerbated for high-dimensional exposures. Caution must be taken in interpreting causal effects when the assumptions needed for causal inference, such as no unmeasured confounding or consistency, cannot be verified. Consequently, any microbiome features identified should be either validated in experimental studies if possible, or more closely scrutinized according to guidelines for evidence of causation. However, even if conditions for causal inference do not hold, our method may provide better recovery of associational microbiome effects as compared to models applied to the pooled data, when there are features impacted by the confounder.

Some have advocated that microbiome data must be treated as compositional (Gloor et al., 2017). Due to the sum to library size constraint, which is not removed by rarefying but rather made constant across all samples, microbiome data technically lie in a simplex space (Aitchison, 1982). One goal of our funded project is to identify microbial features that can be intervened upon to produce a favorable outcome. Hence we analyze count data, not compositional data where it is impossible to alter a feature without changing at least one other so as to retain the same total sum across features. When microbiome features are high dimensional, and in particular there is no dominating feature, the impact of this issue may be minimal. Moreover, microbiome data often exhibit many zeros and the popular centered log-ratio approach for compositional data applies log transformation after adding an arbitrary pseudocount, the choice of which may impact the analysis (Costea et al., 2014). In cases when compositional analysis is preferred, such as when taxa are summarized at the level of genus or higher typically leading to *p* < *n* with a lower prevalence of zeros, our strategy of standardization could be altered in a straightforward way by replacing penalized least squares with a regularized method for compositional covariates (Lin et al., 2014; Shi et al., 2016).

Depending on the underlying biology, the taxonomic structure or phylogeny may be important in the relationship between the microbiome and outcome. If so, higher power may be achieved by using a different penalty that leverages such information. The group LASSO selects groups of features (Yuan and Lin, 2006) and modifications have been developed for microbiome applications incorporating multiple levels of taxonomic hierarchy (Garcia et al., 2014). Other options include a phylogeny-based penalty that penalizes coefficients along a supplied phylogenetic tree (Xian et al., 2018) or a kernel-based penalty incorporating a desired ecological distance (Randolph et al., 2018). To increase power and address the challenge of FDR control, the hierarchical taxonomic structure could be utilized in a multi-stage FDR controlling approach (Hu et al., 2018). Applications of these methods require the taxa assignments and phylogenetic tree, which may be incompletely elucidated for novel microbial species, or measured with error (Golob et al., 2017; Lindgreen et al., 2016).

While simulation studies showed our proposed approach had higher power and better control of FDR at the nominal level compared to other approaches for most scenarios considered, use of the BH procedure with the debiased LASSO and the iterative SIS procedure failed to control FDR for some cases under high dimensionality. Recently, Javanmard and Javadi (2019) showed that the BH procedure may fail to control FDR using the debiased LASSO due to correlation between estimates, but we found little indication of highly correlated estimates in our simulation studies. Correspondingly, applying the Benjamini-Yekutieli adjustment (Benjamini and Yekutieli, 2001) did not result in better FDR control. Instead, it appears our sample sizes were too small to achieve a high enough probability of the sure screening property, leading to relevant features being screened out by the iterative SIS procedure. While additional methodological advancement is needed for valid inference following both variable screening and selection when sample sizes are small, our method performed competitively in recovering relevant features.

## 7 Conclusion

We have addressed the problem of selecting microbiome features relevant to an outcome of interest under confounding by a categorical variable. Our results indicate that standardization enables more accurate identification of individual microbiome features with an effect on the outcome of interest compared to other variable selection and estimation procedures.

## Supporting information

Supplemental results

## Declarations

### Ethics approval and consent to participate

Not applicable.

### Consent for publication

Not applicable.

### Availability of data and materials

R and R markdown code for simulation studies and data analysis are provided in Additional file 2 — R Code. Sequence data can be found in the NCBI SRA submission library under the following accession numbers: sequencing project IDs #1095844, #1095845, #1095846; SRA identifier #SRP165130.

## Acknowledgements

The authors would like to thank Chaohui Yuan for assistance with R code.

## Authors’ contributions

EG, CW, ZH and PL contributed to the methods development and simulation study design. DPS performed field experiments. AMS and DC performed laboratory experiments. EG performed simulation studies and analyzed results. EG wrote the paper with contributions from CW, ZH, AMS, ST, DPS, and PL.

## Additional Files

Additional file 1 — Supplementary Material

Additional simulation results and data analysis.

Additional file 2 — R Code

R and R markdown code for all simulation studies and data analysis.

