## Supplemental results for "Feature selection and causal analysis for microbiome studies in the presence of confounding using standardization"

Emily Goren

November 25, 2019

### S1 Additional simulation results

Table S1: Simulation results: true positive rate (TPR) and false positive rate (FPR) for identification of population coefficients for $n = 100$ and Poisson features, reported as means over 100 simulation replications. For the debiased LASSO with iterative SIS and BH procedure (labeled "iterSIS-dbLASSO-BH"), FDR control was at the 0.05 level.

| | | Variable Selection Method | | | | | | | | | |
|---|---|---|---|---|---|---|---|---|---|---|---|
| | | iterSIS-dbLASSO-BH | | iterSIS-LASSO | | iterSIS-SCAD | | LASSO | | SCAD | |
| Simulation Scenario | Model | TPR | FPR | TPR | FPR | TPR | FPR | TPR | FPR | TPR | FPR |
| Strong Effect Modifier, $p = 50$, 0% confounded | Conditional Std | **1.00** | 0.02 | **1.00** | **0.05** | **1.00** | **0.02** | **1.00** | **0.02** | **1.00** | **0.00** |
| | Select L | 0.04 | 0.04 | 0.96 | 0.96 | 0.87 | 0.80 | 0.96 | 0.96 | 0.87 | 0.80 |
| | Select L EffMod | 0.96 | **0.00** | 0.96 | 0.08 | 0.97 | 0.05 | **1.00** | 0.03 | **1.00** | **0.00** |
| | Require L | 0.03 | 0.03 | 0.95 | 0.95 | 0.86 | 0.81 | 0.95 | 0.95 | 0.86 | 0.81 |
| | Require L EffMod | 0.96 | **0.00** | 0.96 | 0.08 | 0.97 | 0.05 | **1.00** | **0.02** | **1.00** | **0.00** |
| | Ignore L | 0.09 | 0.09 | 0.96 | 0.95 | 0.86 | 0.80 | 0.96 | 0.95 | 0.87 | 0.80 |
| | Ignore L EffMod | 0.97 | **0.00** | 0.97 | 0.08 | 0.97 | 0.04 | **1.00** | **0.02** | **1.00** | **0.00** |
| Strong Effect Modifier, $p = 50$, 60% confounded | Conditional Std | **1.00** | 0.01 | **1.00** | **0.04** | **0.99** | **0.02** | **1.00** | **0.02** | **1.00** | **0.00** |
| | Select L | 0.18 | 0.05 | 0.97 | 0.95 | 0.81 | 0.79 | 0.97 | 0.95 | 0.81 | 0.79 |
| | Select L EffMod | 0.90 | **0.00** | 0.95 | 0.09 | 0.96 | 0.05 | **1.00** | 0.03 | **1.00** | **0.00** |
| | Require L | 0.13 | 0.03 | 0.95 | 0.94 | 0.81 | 0.79 | 0.95 | 0.94 | 0.81 | 0.79 |
| | Require L EffMod | 0.89 | **0.00** | 0.96 | 0.08 | 0.96 | 0.04 | **1.00** | **0.02** | **1.00** | **0.00** |
| | Ignore L | 0.29 | 0.10 | 0.95 | 0.94 | 0.82 | 0.79 | 0.95 | 0.94 | 0.82 | 0.79 |
| | Ignore L EffMod | 0.90 | **0.00** | 0.96 | 0.09 | 0.97 | 0.05 | **1.00** | **0.02** | **1.00** | **0.00** |
| Strong Effect Modifier, $p = 50$, 100% confounded | Conditional Std | **1.00** | 0.01 | **1.00** | **0.02** | **1.00** | **0.00** | **1.00** | **0.02** | **1.00** | **0.00** |
| | Select L | 0.32 | 0.04 | 0.94 | 0.91 | 0.81 | 0.71 | 0.95 | 0.91 | 0.81 | 0.71 |
| | Select L EffMod | 0.91 | **0.00** | 0.99 | 0.06 | **1.00** | 0.03 | **1.00** | **0.02** | **1.00** | **0.00** |
| | Require L | 0.24 | 0.02 | 0.93 | 0.89 | 0.81 | 0.72 | 0.93 | 0.89 | 0.81 | 0.72 |
| | Require L EffMod | 0.92 | **0.00** | 0.99 | 0.06 | **1.00** | 0.02 | **1.00** | **0.02** | **1.00** | **0.00** |
| | Ignore L | 0.44 | 0.07 | 0.92 | 0.89 | 0.81 | 0.71 | 0.92 | 0.89 | 0.81 | 0.71 |
| | Ignore L EffMod | 0.92 | **0.00** | 0.99 | 0.06 | **1.00** | 0.02 | **1.00** | **0.02** | **1.00** | **0.00** |
| Strong Effect Modifier, $p = 2000$, 0% confounded | Conditional Std | **0.55** | **0.00** | **0.61** | **0.01** | **0.63** | **0.01** | **1.00** | **0.01** | **1.00** | **0.00** |
| | Select L | 0.00 | **0.00** | 0.00 | **0.01** | 0.00 | **0.01** | 0.05 | 0.05 | 0.02 | 0.02 |
| | Select L EffMod | 0.01 | **0.00** | 0.01 | **0.01** | 0.01 | **0.01** | **1.00** | **0.01** | **1.00** | **0.00** |
| | Require L | 0.00 | **0.00** | 0.00 | **0.01** | 0.00 | **0.01** | 0.05 | 0.05 | 0.02 | 0.02 |
| | Require L EffMod | 0.01 | **0.00** | 0.01 | **0.01** | 0.00 | **0.01** | **1.00** | **0.01** | **1.00** | **0.00** |
| | Ignore L | 0.00 | 0.01 | 0.00 | **0.01** | 0.00 | **0.01** | 0.05 | 0.05 | 0.02 | 0.02 |
| | Ignore L EffMod | 0.00 | **0.00** | 0.00 | **0.01** | 0.01 | **0.01** | **1.00** | **0.01** | **1.00** | **0.00** |
| Strong Effect Modifier, $p = 2000$, 60% confounded | Conditional Std | **0.48** | **0.00** | **0.52** | **0.01** | **0.54** | **0.01** | **0.99** | **0.01** | **1.00** | **0.00** |
| | Select L | 0.06 | **0.00** | 0.09 | **0.01** | 0.09 | **0.01** | 0.09 | 0.05 | 0.07 | 0.02 |
| | Select L EffMod | 0.11 | **0.00** | 0.15 | **0.01** | 0.17 | **0.01** | **0.99** | **0.01** | **1.00** | **0.00** |
| | Require L | 0.01 | **0.00** | 0.09 | **0.01** | 0.09 | **0.01** | 0.10 | 0.04 | 0.06 | 0.02 |
| | Require L EffMod | 0.13 | **0.00** | 0.15 | **0.01** | 0.17 | **0.01** | **0.99** | **0.01** | **1.00** | **0.00** |
| | Ignore L | 0.07 | 0.01 | 0.09 | **0.01** | 0.10 | **0.01** | 0.09 | 0.05 | 0.07 | 0.02 |
| | Ignore L EffMod | 0.12 | **0.00** | 0.16 | **0.01** | 0.17 | **0.01** | **0.99** | **0.01** | **1.00** | **0.00** |
| Strong Effect Modifier, $p = 2000$, 100% confounded | Conditional Std | **1.00** | **0.00** | **1.00** | **0.00** | **1.00** | **0.00** | **0.98** | **0.01** | **1.00** | **0.00** |
| | Select L | 0.16 | **0.00** | 0.20 | 0.01 | 0.24 | 0.01 | 0.17 | 0.04 | 0.11 | 0.02 |
| | Select L EffMod | 0.30 | **0.00** | 0.36 | 0.01 | 0.38 | 0.01 | **0.98** | **0.01** | **1.00** | **0.00** |
| | Require L | 0.03 | **0.00** | 0.22 | 0.01 | 0.24 | 0.01 | 0.17 | 0.04 | 0.11 | 0.02 |
| | Require L EffMod | 0.32 | **0.00** | 0.36 | 0.01 | 0.37 | 0.01 | **0.98** | **0.01** | **1.00** | **0.00** |
| | Ignore L | 0.19 | 0.01 | 0.22 | 0.01 | 0.25 | 0.01 | 0.17 | 0.04 | 0.11 | 0.02 |
| | Ignore L EffMod | 0.31 | **0.00** | 0.37 | 0.01 | 0.39 | 0.01 | **0.98** | **0.01** | **1.00** | **0.00** |
| No Effect Modifier, $p = 50$, 0% confounded | Conditional Std | **1.00** | 0.02 | **1.00** | 0.05 | **1.00** | **0.02** | **1.00** | 0.02 | **1.00** | **0.00** |
| | Select L | **1.00** | 0.06 | **1.00** | 0.14 | **1.00** | 0.05 | **1.00** | 0.14 | **1.00** | 0.05 |
| | Select L EffMod | 0.97 | **0.00** | 0.97 | 0.08 | 0.97 | 0.04 | **1.00** | 0.03 | **1.00** | **0.00** |
| | Require L | **1.00** | **0.00** | **1.00** | **0.00** | **1.00** | 0.04 | **1.00** | **0.00** | **1.00** | 0.04 |
| | Require L EffMod | 0.97 | **0.00** | 0.97 | 0.08 | 0.97 | 0.04 | **1.00** | 0.02 | **1.00** | **0.00** |
| | Ignore L | **1.00** | 0.11 | **1.00** | 0.14 | **1.00** | 0.05 | **1.00** | 0.14 | **1.00** | 0.05 |
| | Ignore L EffMod | 0.97 | **0.00** | 0.97 | 0.09 | 0.97 | 0.04 | **1.00** | 0.02 | **1.00** | **0.00** |
| No Effect Modifier, $p = 50$, 60% confounded | Conditional Std | **1.00** | 0.01 | **1.00** | 0.04 | 0.99 | **0.02** | **1.00** | **0.02** | **1.00** | **0.00** |
| | Select L | 0.99 | 0.05 | 0.99 | 0.13 | **1.00** | 0.03 | 0.99 | 0.13 | **1.00** | 0.03 |
| | Select L EffMod | 0.77 | **0.00** | 0.96 | 0.09 | 0.96 | 0.05 | **1.00** | **0.02** | **1.00** | **0.00** |
| | Require L | 0.97 | **0.00** | 0.97 | **0.03** | **1.00** | 0.03 | 0.97 | 0.03 | **1.00** | 0.03 |
| | Require L EffMod | 0.77 | **0.00** | 0.96 | 0.08 | 0.96 | 0.05 | **1.00** | **0.02** | **1.00** | **0.00** |
| | Ignore L | 0.99 | 0.07 | 0.99 | 0.12 | **1.00** | 0.03 | 0.99 | 0.12 | **1.00** | 0.03 |
| | Ignore L EffMod | 0.76 | **0.00** | 0.96 | 0.09 | 0.96 | 0.05 | **1.00** | **0.02** | **1.00** | **0.00** |
| No Effect Modifier, $p = 50$, 100% confounded | Conditional Std | **1.00** | 0.02 | **1.00** | **0.02** | **1.00** | **0.00** | **1.00** | **0.02** | **1.00** | **0.00** |
| | Select L | 0.98 | 0.04 | 0.98 | 0.13 | 0.99 | 0.03 | 0.98 | 0.14 | 0.99 | 0.03 |
| | Select L EffMod | 0.70 | 0.01 | **1.00** | 0.07 | **1.00** | 0.03 | **1.00** | 0.03 | 0.99 | **0.00** |
| | Require L | 0.95 | **0.00** | 0.96 | 0.05 | 0.99 | 0.02 | 0.96 | 0.05 | 0.99 | 0.02 |
| | Require L EffMod | 0.70 | **0.00** | **1.00** | 0.06 | **1.00** | 0.02 | **1.00** | **0.02** | **1.00** | **0.00** |
| | Ignore L | 0.98 | 0.05 | 0.98 | 0.13 | 0.99 | 0.03 | 0.98 | 0.13 | 0.99 | 0.03 |
| | Ignore L EffMod | 0.69 | **0.00** | **1.00** | 0.06 | **1.00** | 0.02 | **1.00** | **0.02** | **1.00** | **0.00** |
| No Effect Modifier, $p = 2000$, 0% confounded | Conditional Std | 0.55 | **0.00** | **0.61** | **0.01** | 0.63 | **0.01** | **1.00** | 0.01 | **1.00** | **0.00** |
| | Select L | 0.60 | **0.00** | **0.61** | **0.01** | 0.64 | **0.01** | **1.00** | 0.01 | **1.00** | **0.00** |
| | Select L EffMod | 0.01 | **0.00** | 0.01 | **0.01** | 0.01 | **0.01** | **1.00** | 0.01 | **1.00** | **0.00** |
| | Require L | 0.40 | **0.00** | 0.57 | **0.01** | 0.60 | **0.01** | **1.00** | **0.00** | **1.00** | **0.00** |
| | Require L EffMod | 0.01 | **0.00** | 0.01 | **0.01** | 0.01 | **0.01** | **1.00** | 0.01 | **1.00** | **0.00** |
| | Ignore L | **0.61** | **0.00** | **0.61** | **0.01** | **0.65** | **0.01** | **1.00** | 0.01 | **1.00** | **0.00** |
| | Ignore L EffMod | 0.01 | **0.00** | 0.01 | **0.01** | 0.01 | **0.01** | **1.00** | 0.01 | **1.00** | **0.00** |
| No Effect Modifier, $p = 2000$, 60% confounded | Conditional Std | 0.48 | **0.00** | 0.53 | **0.01** | 0.54 | **0.01** | **0.99** | 0.01 | **1.00** | **0.00** |
| | Select L | 0.54 | **0.00** | 0.56 | **0.01** | 0.62 | **0.01** | 0.96 | 0.01 | 0.97 | **0.00** |
| | Select L EffMod | 0.11 | **0.00** | 0.19 | **0.01** | 0.19 | **0.01** | **0.99** | 0.01 | **1.00** | **0.00** |
| | Require L | 0.42 | **0.00** | 0.55 | **0.01** | 0.59 | **0.01** | 0.92 | **0.00** | 0.97 | **0.00** |
| | Require L EffMod | 0.12 | **0.00** | 0.19 | **0.01** | 0.19 | **0.01** | **0.99** | 0.01 | **1.00** | **0.00** |
| | Ignore L | **0.56** | **0.00** | **0.59** | **0.01** | **0.65** | **0.01** | 0.96 | 0.01 | 0.97 | **0.00** |
| | Ignore L EffMod | 0.12 | **0.00** | 0.20 | **0.01** | 0.20 | **0.01** | **0.99** | 0.01 | **1.00** | **0.00** |
| No Effect Modifier, $p = 2000$, 100% confounded | Conditional Std | **1.00** | **0.00** | **1.00** | **0.00** | **1.00** | **0.00** | **0.99** | **0.01** | **1.00** | **0.00** |
| | Select L | 0.75 | **0.00** | 0.76 | 0.01 | 0.82 | **0.00** | 0.91 | **0.01** | 0.92 | **0.00** |
| | Select L EffMod | 0.23 | **0.00** | 0.44 | 0.01 | 0.45 | 0.01 | **0.99** | **0.01** | 0.99 | **0.00** |
| | Require L | 0.65 | **0.00** | 0.73 | **0.00** | 0.78 | **0.00** | 0.83 | **0.01** | 0.94 | **0.00** |
| | Require L EffMod | 0.24 | **0.00** | 0.44 | 0.01 | 0.46 | 0.01 | 0.98 | **0.01** | **1.00** | **0.00** |
| | Ignore L | 0.78 | **0.00** | 0.78 | 0.01 | 0.88 | **0.00** | 0.91 | **0.01** | 0.92 | **0.00** |
| | Ignore L EffMod | 0.23 | **0.00** | 0.44 | 0.01 | 0.45 | 0.01 | 0.98 | **0.01** | **1.00** | **0.00** |

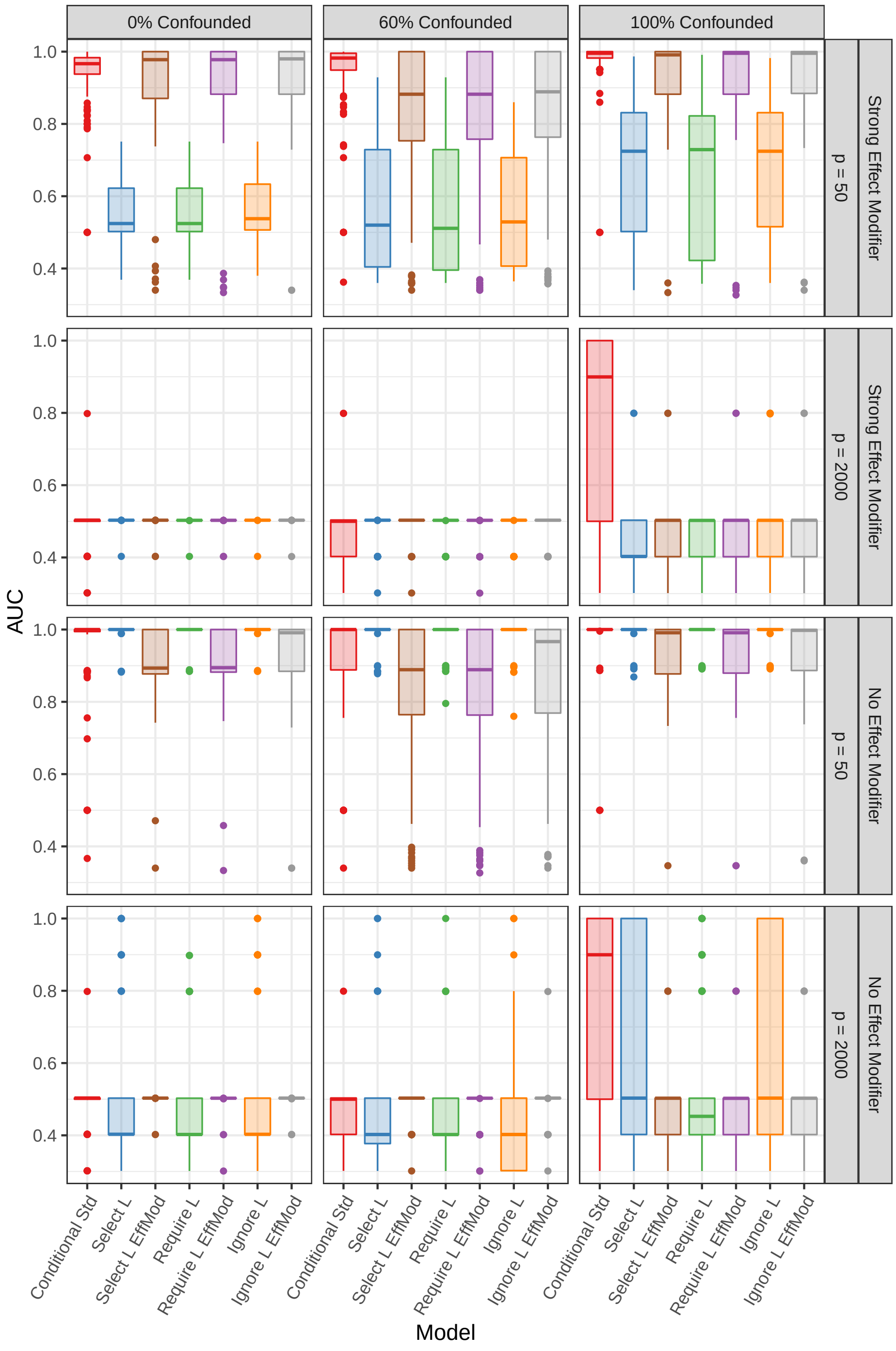

Figure S1: Simulation results: box plots of the area under the curve (AUC) from 100 simulation replications for $n = 50$ and Poisson features using $p$-values based on the debiased LASSO estimate following iterative sure independence screening (iterative SIS).

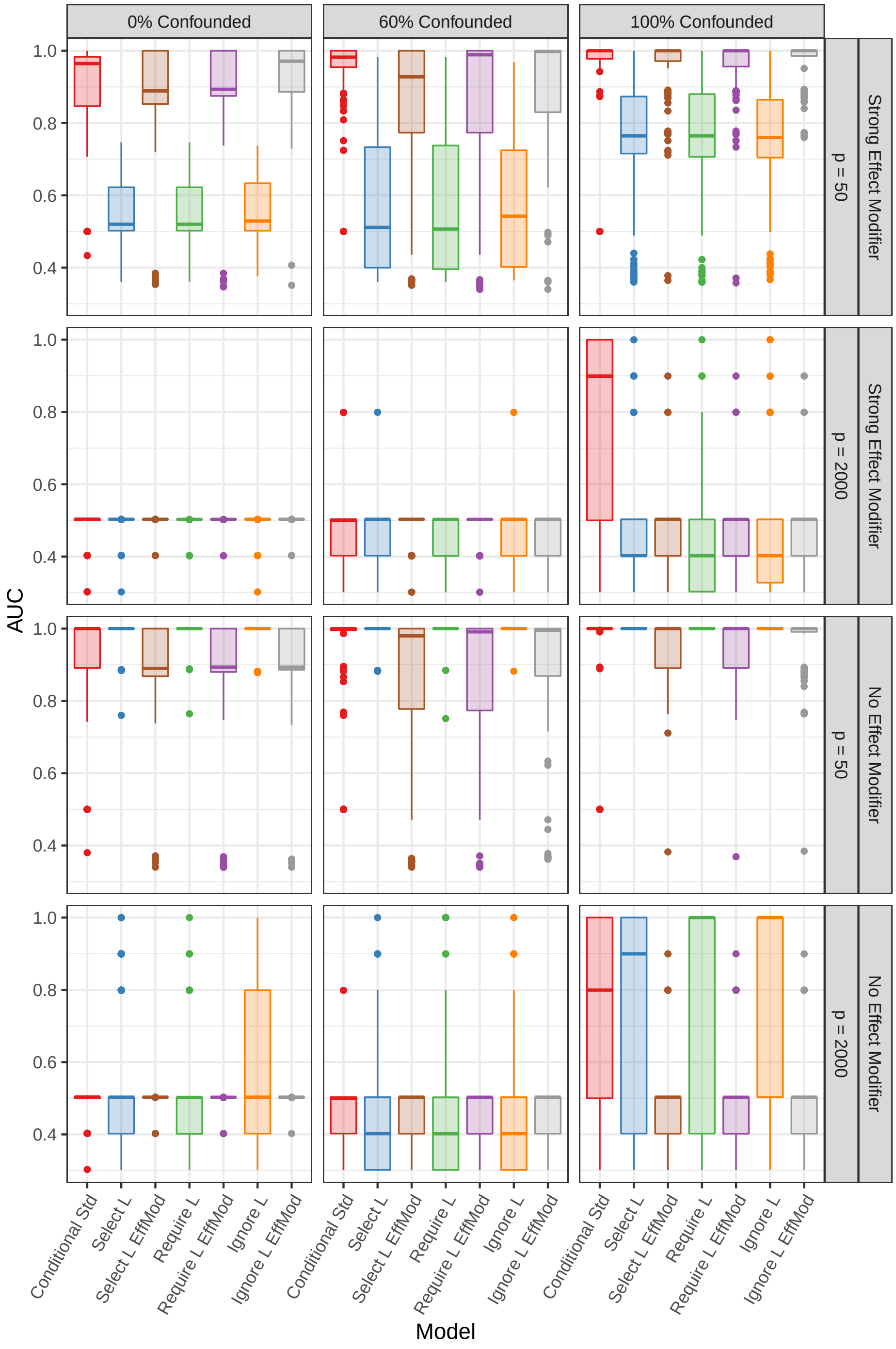

Figure S2: Simulation results: box plots of the area under the curve (AUC) from 100 simulation replications for $n = 50$ and negative binomial features using $p$-values based on the debiased LASSO estimate following iterative sure independence screening (iterative SIS).

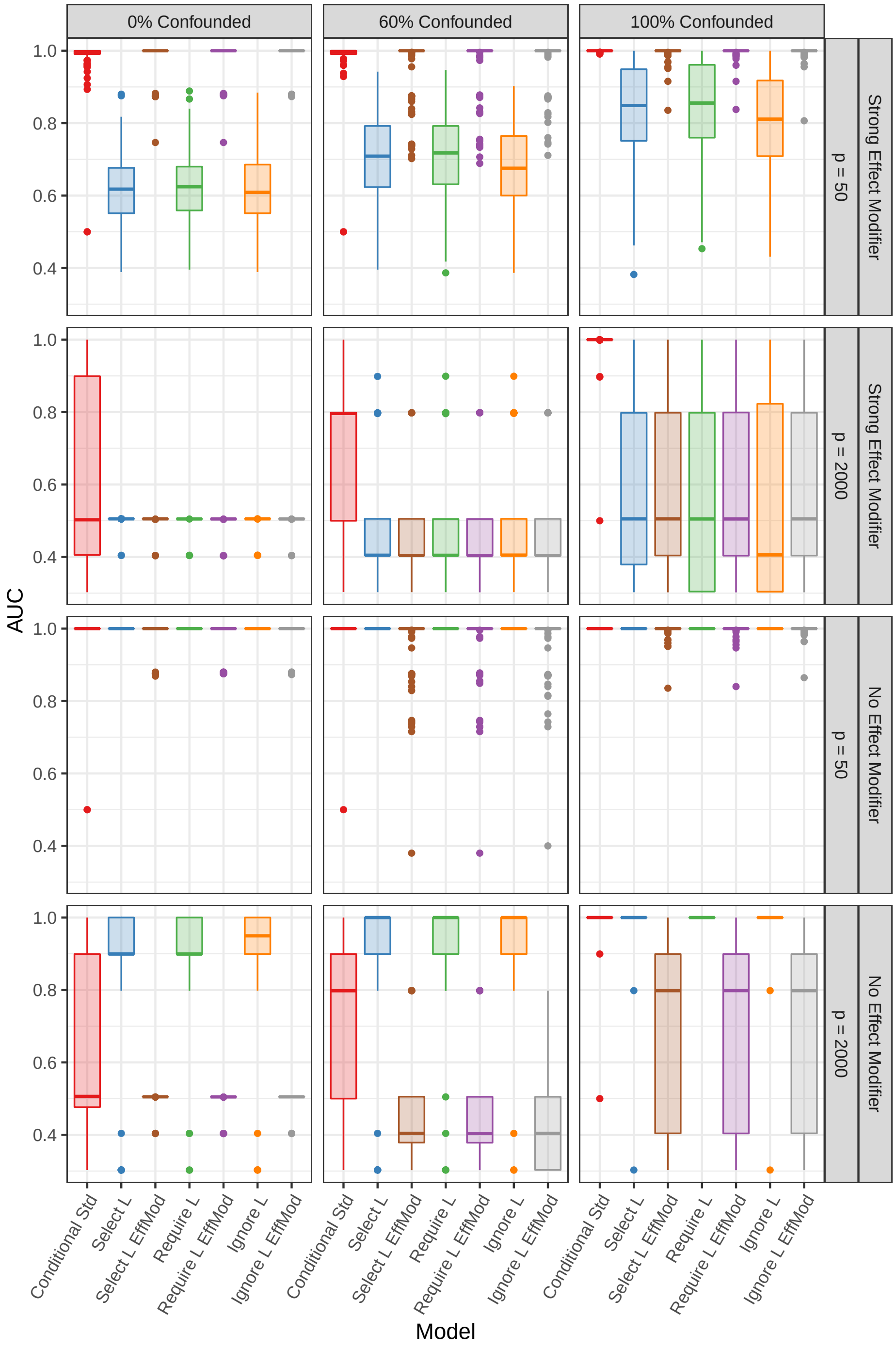

Figure S3: Simulation results: box plots of the area under the curve (AUC) from 100 simulation replications for $n = 100$ and negative binomial features using $p$-values based on the debiased LASSO estimate following iterative sure independence screening (iterative SIS).

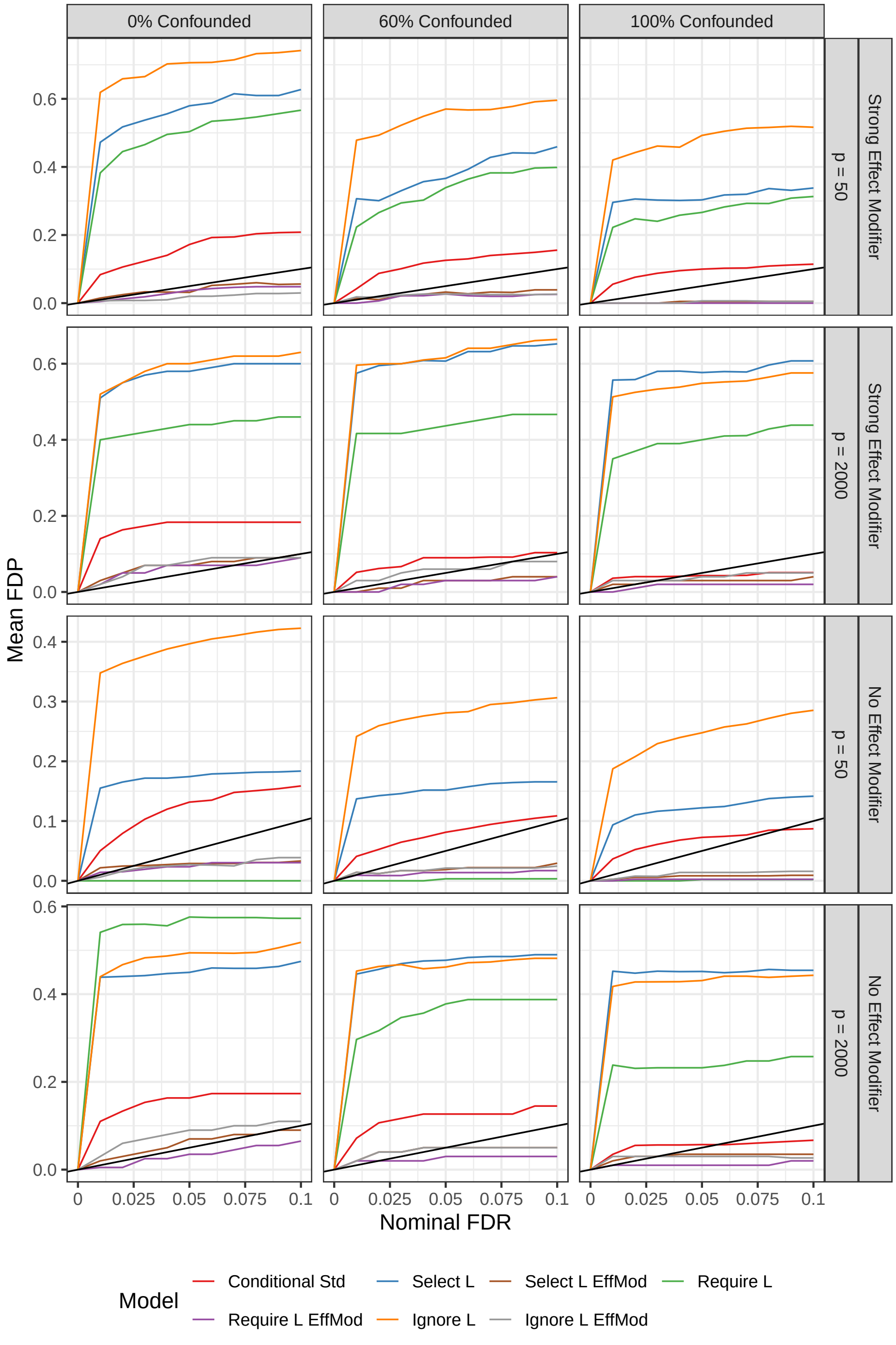

Figure S4: Simulation results: mean estimated false discovery proportion (FDP) for $n = 50$ and Poisson features at varying nominal false discovery rate (FDR) values using Benjamini-Hochberg adjusted $p$-values based on the debiased LASSO estimate following iterative sure independence screening (iterative SIS). The $y = x$ line is shown in black; any values above this line indicate lack of FDR control.

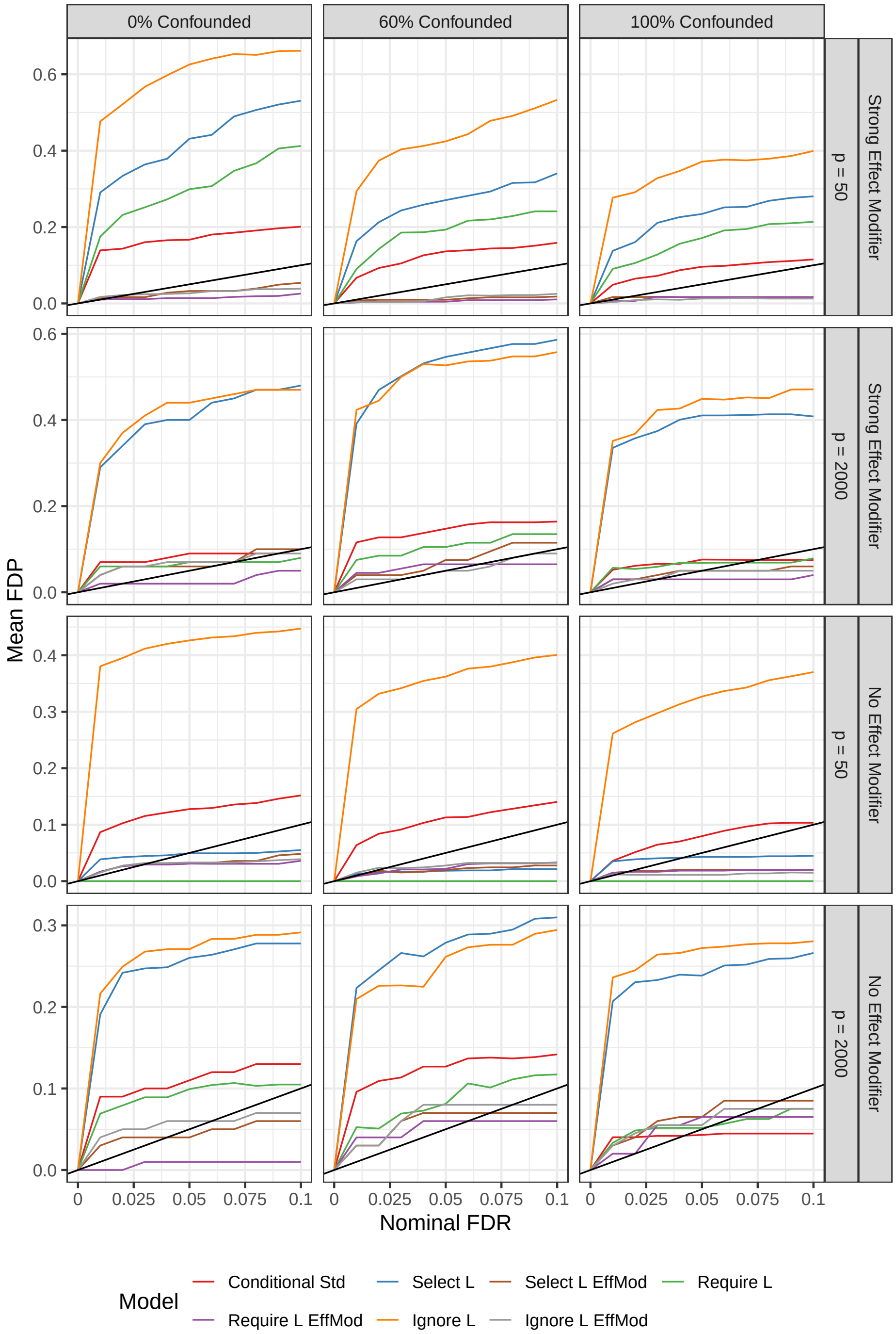

Figure S5: Simulation results: mean estimated false discovery proportion (FDP) for $n = 50$ and negative binomial features at varying nominal false discovery rate (FDR) values using Benjamini-Hochberg adjusted $p$-values based on the debiased LASSO estimate following iterative sure independence screening (iterative SIS). The $y = x$ line is shown in black; any values above this line indicate lack of FDR control.

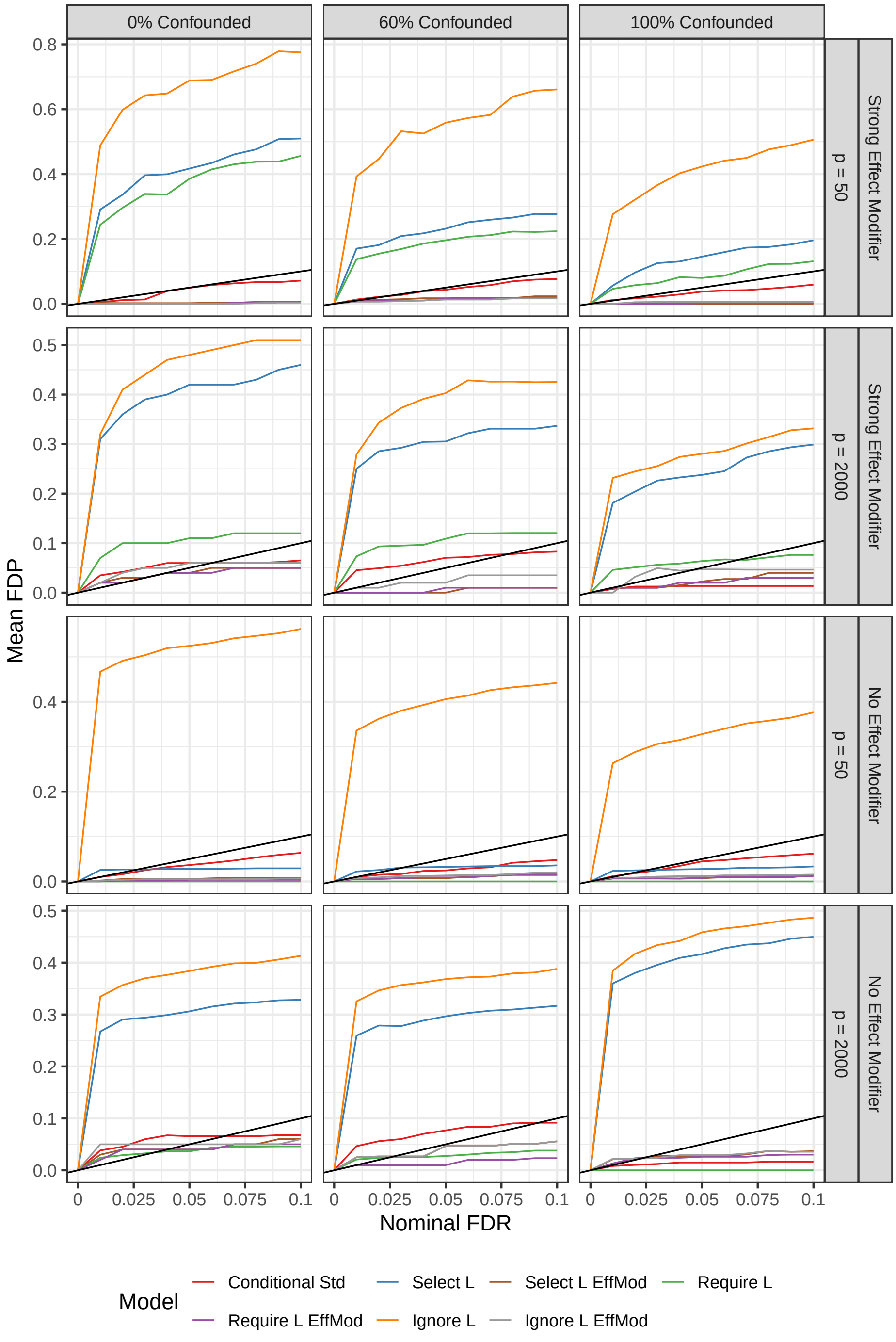

Figure S6: Simulation results: mean estimated false discovery proportion (FDP) for $n = 100$ and negative binomial features at varying nominal false discovery rate (FDR) values using Benjamini-Hochberg adjusted $p$-values based on the debiased LASSO estimate following iterative sure independence screening (iterative SIS). The $y = x$ line is shown in black; any values above this line indicate lack of FDR control.

#### S2 Real data analysis: correlation structure and assumption checks

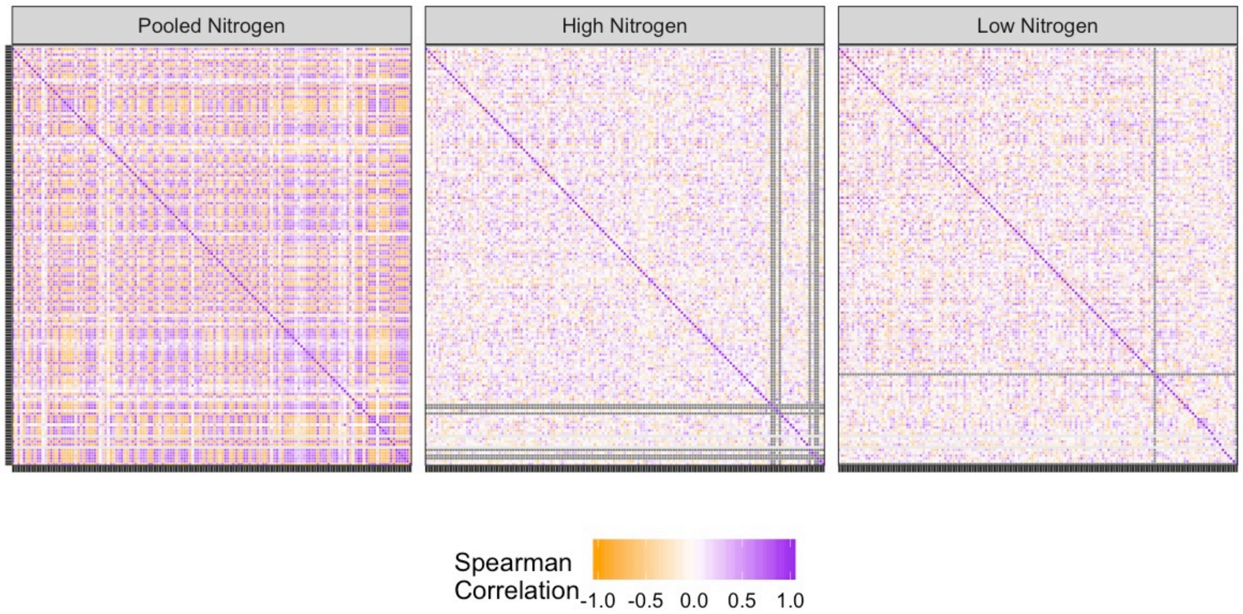

Figure S7: Sorghum microbiome data: Spearman's correlation between the top 150 marginally correlated OTUs for all samples, pooled across both nitrogen conditions (left) and stratified by nitrogen application (center and right).

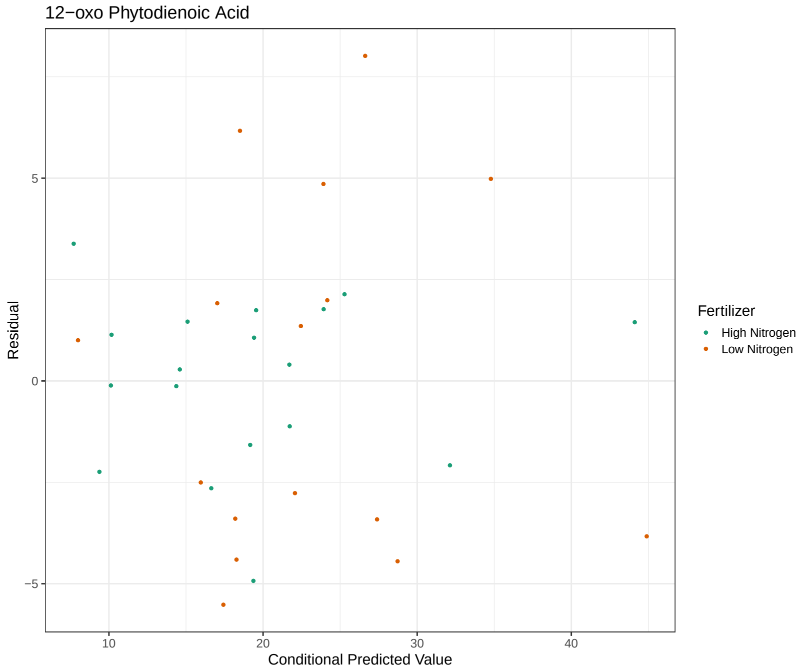

Figure S8: Real data analysis: residuals versus conditional (nitrogen stratum-specific) predicted values from the conditional debiased iterative SIS-LASSO estimates.

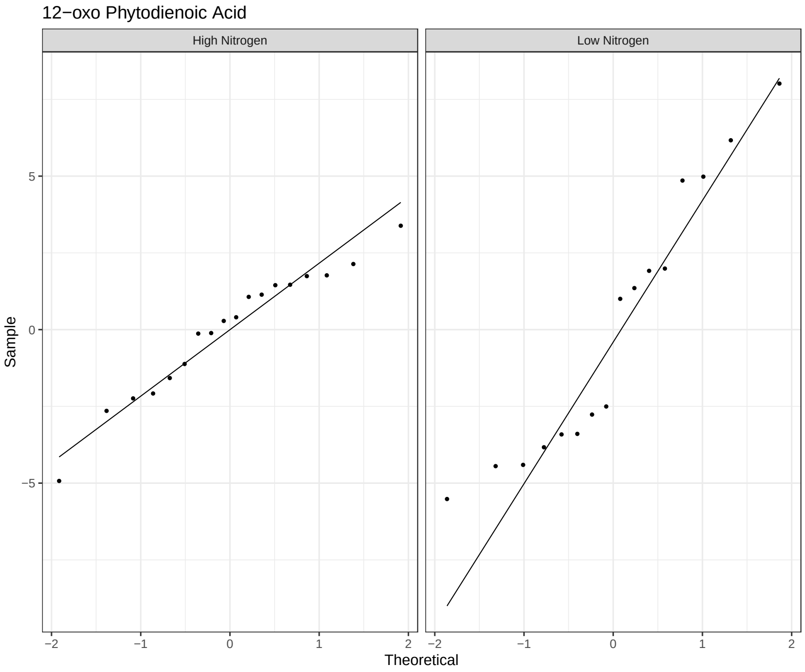

Figure S9: Real data analysis: Q-Q plot of residuals based on conditional (nitrogen stratum-specific) predicted values from the conditional debiased iterative SIS-LASSO estimates.
