## Supplementary material for "Feature selection and causal analysis for microbiome studies in the presence of confounding using standardization": R code: 01_run_simulation.R

```
rm(list = ls())

set.seed(09202018)

require(SISmod)
require(SLOPE)
require(knockoff)
require(hdi)
require(foreach)
require(doParallel)

devtools::session_info()
registerDoParallel(detectCores())


# Number of simulations
N <- 1:100
repIDs <- paste0('rep', N)
# Number of units (observations -- plots, plants, etc)
n <- 100
# Number of OTUs/microbiome features
p <- c(50, 2000)
# Number of OTUs with an effect on response
p.beta <- 5
# Magntude, sign of p.beta, interactions?
betaA <- 3
betaAsign <- 'Mixed'
delta <- c(1, -0.9)
# Treatment effect
betaL <- 5
# Proportion of DA OTUs
prop.DA <- 0.3
# Split of p.beta amoung DA OTUs and nDA OTUs
split <- c(0, 0.5, 1)
# OTUs dsn
dsn <- 'Poisson'
# Standardize X for data generation?
stdA <- TRUE
# Intercept
beta0 <- 3
# Variability (spherical)
sigma <- 0.25
conds <- expand.grid(n = n, p = p, split = split, p.beta = p.beta, 
                     beta0 = beta0, betaA = betaA, betaL = betaL, sigma = sigma,
                     betaAsign = betaAsign, dsn = dsn, stdA = stdA, delta = delta)
J <- nrow(conds)
conds$ID <- 1:J
J

alphas <- 0.05


# Function to generate parameters and data for a given setting.
rdata <- function(params, nsim, ID) {
  set.seed(nsim)
  n <- params$n; p <- params$p; split <- params$split; p.beta <- params$p.beta
  beta0 <- params$beta0; betaA <- params$betaA; betaL <- params$betaL
  sigma <- params$sigma; betaAsign <- params$betaAsign; dsn <- params$dsn; 
  stdA <- params$stdA; delta <- params$delta
  # Indicate if OTU is DA
  is.DA <- c(rep(TRUE, prop.DA*p), rep(FALSE, (1-prop.DA)*p))
  # Indicate treatment/confounder
  L <- rep(c(1,0), each = (n/2))
  ###### Set phi (neg bin dispersion)
  if (dsn == 'Negative Binomial') {
    phi <- rexp(p, rate = 1/5)
  } else {
    phi <- rep(1e6, p)
  }
  ###### Set beta (effect of microbiome on Y)
  # Number of non-zero betas in DA OTUs
  p.DA <- ceiling(split * p.beta)
  # Number of non-zero betas in non-DA OTUs
  p.nDA <- p.beta - p.DA
  idx <- c(if(p.DA > 1) which(is.DA)[1:p.DA], if(p.nDA > 1) which(!is.DA)[1:p.nDA])
  #idx <- c(if(p.DA > 1) (sum(is.DA)-p.DA+1):(sum(is.DA)), if(p.nDA > 1) (p-p.nDA+1):p)
  beta <- rep(0, p)
  if (betaAsign == 'Positive') {
    beta[idx] <- betaA
  } else if (betaAsign == 'Mixed') {
    beta[idx] <- betaA * rep_len(c(1,-1), p.beta)
  } else if (betaAsign == 'Negative') {
    beta[idx] <- (-1 * betaA)
  }
  ##### Set gamma (microbiome params)
  # Intercept for each OTU
  gamma0 <- rlnorm(p, meanlog = 1/2, sdlog = 3/2)
  gamma0[idx] <- 5
  # DA effect for each OTU
  gamma1 <- rep(1, p)
  m <- sample(rep(c(1, -1), each = sum(is.DA)/2))
  gamma1[is.DA] <- rlnorm(sum(is.DA), meanlog = m*(1/4), sdlog = 3/2)
  ##### Draw OTUs
  if (dsn %in% c('Poisson', 'Negative Binomial')) {
    A <- sapply(1:p, function(j) MASS::rnegbin(n, mu = gamma0[j] * (gamma1[j]^L), theta = phi[j]))
  } else {
    gamma0[idx] <- 10
    gamma0[is.DA] <- 10
    gamma1[is.DA] <- 10
    A <- sapply(1:p, function(j) MASS::rnegbin(n, mu = gamma0[j] * (gamma1[j]^L), theta = phi[j]))
  } 
  colnames(A) <- 1:p
  ##### Draw Y
  if (stdA) {
    X <- cbind(SISmod::scale(A), L)
  } else {
    X <- cbind(A, L)
  }
  X[is.na(X)] <- 0
  beta.cond0 <- beta
  beta.cond1 <- delta * beta
  beta.std <- (beta.cond0 + beta.cond1) / 2
  Y0 <- rnorm(sum(L == 0), X[L == 0,] %*% c(beta.cond0, betaL), sigma)
  Y1 <- rnorm(sum(L == 1), X[L == 1,] %*% c(beta.cond1, betaL), sigma)
  Y <- c(Y1, Y0)
  # Output.
  out <- list(L, A, Y, beta.std, params, nsim)
  names(out) <- c('L', 'A', 'Y', 'beta', 'params', 'nsim')
  return(out)
}

# SIS function.
sis <- function(x, y, d) {
    cors <- suppressWarnings(apply(x, 2, cor, y = y))
    cors[is.na(cors)] <- 0
    ans <- order(abs(cors), decreasing = TRUE)[1:d]
    return(ans)
}

# Function to extract selected coef indices
get.sel <- function(fit, alpha) {
    switch(class(fit)[1],
           logical = NA,
           numeric = names(which(fit < alpha)),
           hdi = names(which(fit$pval.corr < alpha)),
           knockoff.result = names(fit$selected),
           SLOPE.result = names(fit$selected),
           list = {
               sl <- names(fit$ix)
               if (is.null(sl)) sl <- fit$ix
               return(sl)
           })
}

# Functions to extract selected results.
sumdat <- function(res, alphas, p, std = FALSE) {
    nms <- names(res)
    ans <- lapply(1:length(res), function(r) {
        if (nms[r] %in% c('ISIS-Slope', 'ISIS-Knockoff', 'SIS-SLOPE', 'SIS-Knockoff')) {
            df <- data.frame(sel = t(sapply(1:length(alphas), function(a) {
                if (std) {
                    sl <- get.sel(res[[r]][[a]], alphas[a])
                    sl0 <- (1:p) %in% sl
                    sl1 <- ((p+1):(2*p)) %in% sl
                    ans <- (sl0 + sl1) > 0
                } else {
                    ans <- (1:p) %in% get.sel(res[[r]][[a]], alphas[a])
                }
                return(ans) })), 
                Method = nms[r], alpha = alphas)
        } else if (nms[r] %in% c('db-SIS-LASSO-BH', 'db-SISLASSO-BY', 'db-SISLASSO-bootWY',
                                 'db-ISIS-LASSO-BH', 'db-ISIS-LASSO-BY', 'db-ISIS-LASSO-bootWY')) {
            df <- data.frame(sel = t(sapply(1:length(alphas), function(a) {
                if (std) {
                    sl <- get.sel(res[[r]], alphas[a])
                    sl0 <- (1:p) %in% sl
                    sl1 <- ((p+1):(2*p)) %in% sl
                    ans <- (sl0 + sl1) > 0
                } else {
                    ans <- (1:p) %in% get.sel(res[[r]], alphas[a])
                }
                return(ans) })), 
                Method = nms[r], alpha = alphas)
        } else {
            if (std) {
                sl <- get.sel(res[[r]], 1)
                sl0 <- (1:p) %in% sl
                sl1 <- ((p+1):(2*p)) %in% sl
                ans <- (sl0 + sl1) > 0
            } else {
                ans <- (1:p) %in% get.sel(res[[r]], 1)
            }
            df <- data.frame(sel = t(ans), Method = nms[r], alpha = NA)
        }
        return(df)
    })
    ans <- do.call('rbind', ans)
    return(ans)
}
sumdatcond <- function(res0, res1, alphas, p) {
    nms <- names(res0)
    ans <- lapply(1:length(res0), function(r) {
        if (nms[r] %in% c('ISIS-Slope', 'ISIS-Knockoff', 'SIS-SLOPE', 'SIS-Knockoff')) {
            df <- data.frame(sel = t(sapply(1:length(alphas), function(a) {
                sl0 <- (1:p) %in% get.sel(res0[[r]][[a]], alphas[a])
                sl1 <- (1:p) %in% get.sel(res1[[r]][[a]], alphas[a])
                ans <- (sl0 + sl1) > 0
                return(ans) })),
                Method = nms[r], alpha = alphas)
        } else if (nms[r] %in% c('db-SIS-LASSO-BH', 'db-SISLASSO-BY', 'db-SISLASSO-bootWY',
                                 'db-ISIS-LASSO-BH', 'db-ISIS-LASSO-BY', 'db-ISIS-LASSO-bootWY')) {
            df <- data.frame(sel = t(sapply(1:length(alphas), function(a) {
                sl0 <- (1:p) %in% get.sel(res0[[r]], alphas[a])
                sl1 <- (1:p) %in% get.sel(res1[[r]], alphas[a])
                ans <- (sl0 + sl1) > 0
                return(ans) })),
                Method = nms[r], alpha = alphas)
        } else {
            sl0 <- (1:p) %in% get.sel(res0[[r]], 1)
            sl1 <- (1:p) %in% get.sel(res1[[r]], 1)
            ans <- (sl0 + sl1) > 0
            df <- data.frame(sel = t(ans), Method = nms[r], alpha = NA)
        }
        return(df)
    })
    ans <- do.call('rbind', ans)
    return(ans)
}

# Function to fits models given x and y.
# For keepL = TRUE, L must correspond to last column of x.
fit <- function(x, y, keepL = FALSE, tune = 'bic', std = FALSE, alphas) {
    yc <- (y - mean(y))
    n <- length(y); p <- ncol(x)
    if (colnames(x)[p] == '') colnames(x)[p] <- p
    penalty.factor <- rep(1, p)
    keep.vars <- rep(FALSE, p)
    d <- floor(n/log(n))
    SIS.idx <- sis(x, y, d)
    if (keepL) {
        penalty.factor[p] <- 0
        keep.vars[p] <- TRUE
        if (!(p %in% SIS.idx)) SIS.idx <- c(SIS.idx, p)
    }
    # LASSO
    lasso <- tryCatch(suppressWarnings(tune.fit(x = x, y = y, penalty = 'lasso', tune = tune, nfolds = n, 
                                                penalty.factor = penalty.factor, standardize = std)),
                      error = function(e) NA)
    ISISlasso <- tryCatch(suppressWarnings(SIS(x = x, y = y, penalty = 'lasso', tune = tune, nfolds = n, 
                                               keep.vars = keep.vars, standardize = std)),
                          error = function(e) NA)
    SISlasso <- tryCatch(suppressWarnings(tune.fit(x = x[,SIS.idx], y = y, penalty = 'lasso', tune = tune, nfolds = n, 
                                                   penalty.factor = penalty.factor[SIS.idx], standardize = std)),
                         error = function(e) NA)
    b.SISlasso <- SISlasso$beta
    x.SISlasso <- x[,as.numeric(names(b.SISlasso))]
    b.ISISlasso <- ISISlasso$coef.est[-1]
    idx <- as.numeric(sapply(names(b.ISISlasso), function(s) gsub('X', '', s)))
    x.ISISlasso <- x[,idx]
    # SCAD
    scad <- tryCatch(suppressWarnings(tune.fit(x = x, y = y, penalty = 'SCAD', tune = tune, nfolds = n, 
                                               penalty.factor = penalty.factor, standardize = std)),
                     error = function(e) NA)
    ISISscad <- tryCatch(suppressWarnings(SIS(x = x, y = y, penalty = 'SCAD', tune = tune, nfolds = n, 
                                              keep.vars = keep.vars, standardize = std)),
                         error = function(e) NA)
    SISscad <- tryCatch(suppressWarnings(tune.fit(x = x[,SIS.idx], y = y, penalty = 'SCAD', tune = tune, nfolds = n, penalty.factor = penalty.factor[SIS.idx], standardize = std)),
                        error = function(e) NA)
    # SLOPE
    SISslope <- lapply(alphas, function(alpha) {
        if (keepL) {
            SIS.lams <- c(create_lambda(n, d, fdr = alpha, method = 'bhq'), 0)
        } else {
            SIS.lams <- create_lambda(n, d, fdr = alpha, method = 'bhq')
        }
        out <- tryCatch(SLOPE(X = x[,SIS.idx], y = yc, fdr = alpha, lambda = SIS.lams, normalize = FALSE), 
                        error = function(e) NA)
        return(out)
    })
    
    # knockoff
    SISknockoff <- lapply(alphas, function(alpha) {
        tryCatch(knockoff.filter(X = x[,SIS.idx], y = y, knockoffs = create.fixed, fdr = alpha),
                 error = function(e) NA)
    })
    names(SISknockoff) <- names(SISslope) <- alphas
    
    # de-biased LASSO tests with FDR adjustment following screening
    if (class(x.SISlasso) != 'matrix') x.SISlasso <- matrix(x.SISlasso, ncol = 1)
    sigma.SISlasso <- sqrt(sum((yc - x.SISlasso%*%b.SISlasso)^2) / (n - length(b.SISlasso) - 1)) # naive est of sigma
    SISlasso.bh <- tryCatch(suppressWarnings(lasso.proj(x = x.SISlasso, y = y, standardize = std, suppress.grouptesting = TRUE, sigma = sigma.SISlasso,
                                                        multiplecorr.method = 'BH', betainit = b.SISlasso, return.Z = TRUE)), error = function(e) NA)
    # Bootstrap w/ Westfall-Young FWER
    SISlasso.wy <- tryCatch(suppressWarnings(lasso.proj(x = x.SISlasso, y = y, standardize = std, suppress.grouptesting = TRUE, sigma = sigma.SISlasso,
                                                        multiplecorr.method = 'WY', betainit = b.SISlasso, Z = SISlasso.bh$Z)), error = function(e) NA)
    if (class(x.ISISlasso) != 'matrix') x.ISISlasso <- matrix(x.ISISlasso, ncol = 1)
    sigma.ISISlasso <- sqrt(sum((yc - x.ISISlasso%*%b.ISISlasso)^2) / (n - length(b.ISISlasso) - 1)) # naive est of sigma
    ISISlasso.bh <- tryCatch(suppressWarnings(lasso.proj(x = x.ISISlasso, y = y, standardize = std, suppress.grouptesting = TRUE, sigma = sigma.ISISlasso,
                                                         multiplecorr.method = 'BH', betainit = b.ISISlasso, return.Z = TRUE)), error = function(e) NA)
    # Bootstrap w/ Westfall-Young FWER
    ISISlasso.wy <- tryCatch(suppressWarnings(lasso.proj(x = x.ISISlasso, y = y, standardize = std, suppress.grouptesting = TRUE, sigma = sigma.ISISlasso,
                                                         multiplecorr.method = 'WY', betainit = b.ISISlasso, Z = ISISlasso.bh$Z)), error = function(e) NA)
    
    # "Oracle" p-values
    SISlasso.oracleBH <- p.adjust(sapply(1:p, function(j) {
        if (j %in% as.numeric(names(b.SISlasso))) {
            lse <- lm(yc ~ . - 1, data = data.frame(yc, x.SISlasso))
            pv <- tryCatch(summary(lse)$coefficients[paste0('X', j),4],
                           error = function(e) NA)
        } else {
            lse <- lm(yc ~ .-1, data = data.frame(yc, x.SISlasso, x[,j]))
            pv <- tryCatch(summary(lse)$coefficients['x...j.',4],
                           error = function(e) NA)
        }
        return(pv)
    }), method = 'BH')
    ISISlasso.oracleBH <- p.adjust(sapply(1:p, function(j) {
        if (j %in% idx) {
            lse <- lm(yc ~ . - 1, data = data.frame(yc, x.ISISlasso))
            pv <- tryCatch(summary(lse)$coefficients[paste0('X', j),4],
                           error = function(e) NA)
        } else {
            lse <- lm(yc ~ . - 1, data = data.frame(yc, x.ISISlasso, x[,j]))
            pv <- tryCatch(summary(lse)$coefficients['x...j.',4],
                           error = function(e) NA)
        }
        return(pv)
    }), method = 'BH')
    lasso.idx <- as.numeric(names(lasso$beta))
    x.lasso <- x[, lasso.idx]
    lasso.oracleBH <- p.adjust(sapply(1:p, function(j) {
        if (j %in%  lasso.idx) {
            lse <- lm(yc ~ . -1, data = data.frame(yc, x.lasso))
            pv <- tryCatch(summary(lse)$coefficients[paste0('X', j),4],
                           error = function(e) NA)
        } else {
            lse <- lm(yc ~ . -1, data = data.frame(yc, x.lasso, x[,j]))
            pv <- tryCatch(summary(lse)$coefficients['x...j.',4],
                           error = function(e) NA)
        }
        return(pv)
    }), method = 'BH')
    ISISscad.idx <- as.numeric(sapply(names(ISISscad$coef.est[-1]), function(s) gsub('X', '', s)))
    x.ISISscad <- x[,ISISscad.idx]
    ISISscad.oracleBH <- p.adjust(sapply(1:p, function(j) {
        if (j %in% ISISscad.idx) {
            lse <- lm(yc ~ . - 1, data = data.frame(yc, x.ISISscad))
            pv <- tryCatch(summary(lse)$coefficients[paste0('X', j),4],
                           error = function(e) NA)
        } else {
            lse <- lm(yc ~ . -1, data = data.frame(yc, x.ISISscad, x[,j]))
            pv <- tryCatch(summary(lse)$coefficients['x...j.',4],
                           error = function(e) NA)
        }
        return(pv)
    }), method = 'BH')
    b.SISscad <- SISscad$beta
    x.SISscad <- x[,as.numeric(names(b.SISscad))]
    SISscad.oracleBH <- p.adjust(sapply(1:p, function(j) {
        if (j %in% as.numeric(names(b.SISscad))) {
            lse <- lm(yc ~ . -1, data = data.frame(yc, x.SISscad))
            pv <- tryCatch(summary(lse)$coefficients[paste0('X', j),4],
                           error = function(e) NA)
        } else {
            lse <- lm(yc ~ . - 1, data = data.frame(yc, x.SISscad, x[,j]))
            pv <- tryCatch(summary(lse)$coefficients['x...j.',4],
                           error = function(e) NA)
        }
        return(pv)
    }), method = 'BH')
    scad.idx <- as.numeric(names(scad$beta))
    x.scad <- x[, scad.idx]
    scad.oracleBH <- p.adjust(sapply(1:p, function(j) {
        if (j %in%  scad.idx) {
            lse <- lm(yc ~ .-1, data = data.frame(yc, x.scad))
            pv <- tryCatch(summary(lse)$coefficients[paste0('X', j),4],
                           error = function(e) NA)
        } else {
            lse <- lm(yc ~ .-1, data = data.frame(yc, x.scad, x[,j]))
            pv <- tryCatch(summary(lse)$coefficients['x...j.',4],
                           error = function(e) NA)
            }
        return(pv)
    }), method = 'BH')
    names(lasso.oracleBH) <- names(SISlasso.oracleBH) <- names(ISISlasso.oracleBH) <- colnames(x)
    names(scad.oracleBH) <- names(SISscad.oracleBH) <- names(ISISscad.oracleBH) <- colnames(x)    
    fits <- list(SISslope, SISknockoff,
                 SISlasso.bh, SISlasso.wy,
                 ISISlasso.bh, ISISlasso.wy,
                 lasso.oracleBH, SISlasso.oracleBH, ISISlasso.oracleBH,
                 scad.oracleBH, SISscad.oracleBH, ISISscad.oracleBH,
                 lasso, ISISlasso, SISlasso, 
                 scad, ISISscad, SISscad)
    names(fits) <- c('SIS-SLOPE', 'SIS-Knockoff',
                     'db-SIS-LASSO-BH', 'db-SIS-LASSO-bootWY',
                     'db-ISIS-LASSO-BH', 'db-ISIS-LASSO-bootWY',
                     'LASSO-oracle-BH', 'SIS-LASSO-oracle-BH', 'ISIS-LASSO-oracle-BH', 
                     'SCAD-oracle-BH', 'SIS-SCAD-oracle-BH', 'ISIS-SCAD-oracle-BH', 
                     'LASSO', 'ISIS-LASSO', 'SIS-LASSO', 'SCAD', 'ISIS-SCAD', 'SIS-SCAD')
    return(fits)
}

# Function to extract results.
get.res <- function(fitlist, d) {
    p <- d$params$p
    bt <- d$beta
    if (p != length( bt))
        print(paste0('True beta is wrong dimension for condition ', d$ID, ' and nsim ', d$nsim))
    tr <- (bt != 0)[1:p]
    varnames <- paste0('sel.', 1:p)
    fitlist$nsel <- apply(fitlist[, varnames], 1, sum)
    fitlist$FDP <- apply(fitlist[, varnames], 1, function(b) {
        den <- sum(b)
        fdp <- ifelse(den == 0, 0, (b %*% (!tr)) /  den)
        return(fdp)
    })
    fitlist$TP <- apply(fitlist[,varnames], 1, function(b) (b %*% tr))
    fitlist$FP <- apply(fitlist[,varnames], 1, function(b) (b %*% (!tr)))
    fitlist$FNR <- apply(fitlist[, varnames], 1, function(b) ((!b) %*% tr) / sum(tr) )
    fitlist$FPR <- apply(fitlist[, varnames], 1, function(b) (b %*% (!tr)) / sum(!tr) )
    fitlist$TPR <- apply(fitlist[, varnames], 1, function(b) (b %*% tr) / sum(tr)) # = sensitivity/power
    fitlist$TNR <- apply(fitlist[, varnames], 1, function(b) ((!b) %*% (!tr)) / sum(!tr) ) # = specificity
    fitlist$ID <- d$params$ID
    out <- merge(fitlist, d$params, by = 'ID')
    out <- out[, !(names(out) %in% varnames)]
    return(out)
}

# Simulate data and parameters.
foreach(j = 1:J) %dopar% {
  out <- foreach(nsim = N, combine = c) %do% rdata(conds[j,], nsim, j)
  names(out) <- repIDs
  save(out, file = paste0('./dump/data_params_ID', j, '.Rdat'))
  NULL
}

# Fit models.
mods <- c('Model1', 'Model1.int', 'Model2', 
          'Model2.keepL', 'Model2.int', 'Model2.int.keepL',
          'Model3.0', 'Model3.1')
fts <- foreach(j = 1:J, .combine = c) %dopar% {
    load(paste0('./dump/data_params_ID', j, '.Rdat'))
    data <- out
    if (data[[repIDs[1]]]$params$ID != j) print('Condition ID does not match index')
    ft <- foreach(repID = repIDs, .combine = c) %do% {
        print(paste('Fitting models for condition ID', j, 'simulation', repID))
        d <- data[[repID]]
        A1 <- rbind(SISmod::scale(d$A[d$L == 1,]), matrix(0, ncol = ncol(d$A), nrow = sum(d$L == 1)))
        A0 <- rbind(matrix(0, ncol = ncol(d$A), nrow = sum(d$L == 0)), SISmod::scale(d$A[d$L == 0,]))
        X <- cbind(A1, A0)
        out <- foreach(mod = mods, .combine = c) %do% {
            print(paste0('Fitting ', mod, ' for ', repID, ' condition ', j))
            res <- switch(mod,
                          Model1 = fit(x = SISmod::scale(d$A), y = d$Y, alphas = alphas),
                          Model1.int = fit(x = X, y = d$Y, alphas = alphas),
                          Model2 = fit(x = cbind(SISmod::scale(d$A), d$L), y = d$Y, alphas = alphas),
                          Model2.keepL = fit(x = cbind(SISmod::scale(d$A), d$L), y = d$Y, keepL = TRUE, alphas = alphas),
                          Model2.int = fit(x = cbind(X, d$L), y = d$Y, alphas = alphas),
                          Model2.int.keepL = fit(x = cbind(X, d$L), y = d$Y, keepL = TRUE, alphas = alphas),
                          Model3.0 = fit(x = SISmod::scale(d$A[d$L == 0, ]), y = d$Y[d$L == 0], alphas = alphas),
                          Model3.1 = fit(x = SISmod::scale(d$A[d$L == 1, ]), y = d$Y[d$L == 1], alphas = alphas))
            list(res)
        }
        names(out) <- mods
        save(out, file = paste0('./dump/sim_fits_ID', j, '_', repID, '.Rdat'))
        list(out)
    }
    names(ft) <- repIDs
    save(ft, file = paste0('./dump/sim_fits_ID', j, 'all.Rdat'))
    ft
}


# Extract results for model selection.
o <- foreach(j = 1:J, .combine = c) %dopar% {
    load(paste0('./dump/data_params_ID', j, '.Rdat'))
    data <- out
    if (data[[repIDs[1]]]$params$ID != j) print('Condition ID does not match index')
    load(paste0('./dump/sim_fits_ID', j, 'all.Rdat'))
    oj <- foreach(repID = repIDs, .combine = c) %do% {
        print(paste('Finding selected OTUs for condition ID', j, 'simulation', repID))
        thisft <- ft[[repID]]
        d <- data[[repID]]
        p <- d$params$p
        b <- rep(FALSE, p)
        # Model 1: ignore treatment
        o1 <- sumdat(thisft$Model1, alphas = alphas, p = p, std = FALSE)
        o1$Model <- 'Ignore L'
        o1.int.std <- sumdat(thisft$Model1.int, alphas = alphas, p = p, std = TRUE)
        o1.int.std$Model <- 'Ignore L Std'
        # Model 2: additive treatment
        o2 <- sumdat(thisft$Model2, alphas = alphas, p = p, std = FALSE)
        o2$Model <- 'Additive L'
        o2.keepL <- sumdat(thisft$Model2.keepL, alphas = alphas, p = p, std = FALSE)
        o2.keepL$Model <- 'Additive Unpenalized L'
        o2.int.std <- sumdat(thisft$Model2.int, alphas = alphas, p = p, std = TRUE)
        o2.int.std$Model <- 'Additive L Std'
        o2.int.keepL.std <- sumdat(thisft$Model2.int.keepL, alphas = alphas, p = p, std = TRUE)
        o2.int.keepL.std$Model <- 'Additive Unpenalized L Std'
        # Model 3: conditional on treatment
        o3 <- sumdatcond(thisft$Model3.0, thisft$Model3.1, alphas = alphas, p = p)
        o3$Model <- 'Conditional Std'
        out <- list(Ignore = o1, Additive = o2, Additive.keepL = o2.keepL,
                    Ignore.int.std = o1.int.std, Additive.int.std = o2.int.std, 
                    Additive.int.keepL.std = o2.int.keepL.std, Conditional.std = o3)
        out <- do.call('rbind', out)
        out$ID <- d$params$ID
        list(out)
    }
    names(oj) <- repIDs
    list(oj)
}

# Extract and save results in a dataframe.
res <- foreach(j = 1:J, .combine = rbind) %dopar% {
    load(paste0('./dump/data_params_ID', j, '.Rdat'))
    data <- out
    if (data[[repIDs[1]]]$params$ID != j) print('Condition ID does not match index')
    resj <- foreach(repID = repIDs, .combine = rbind) %do% get.res(o[[j]][[repID]], data[[repID]])
    resj$ID <- j
    write.csv(resj, file = paste0('./dump/results_ID', j, '.csv'), row.names = FALSE)
    resj
}
write.csv(res, file = './results.csv', row.names = FALSE)
```
