## Supplementary material for "Feature selection and causal analysis for microbiome studies in the presence of confounding using standardization": R code: 02_simulation_analysis.R

```
rm(list = ls())

require(tidyverse)
require(doParallel)
require(ggsci)

devtools::session_info()
theme_set(theme_bw(base_size = 13))
resall <- read.csv('./results.csv')
rmv <- c('ISIS-LASSO-oracle-BH', 'ISIS-SCAD-oracle-BH',
         'LASSO-oracle-BH', 'SCAD-oracle-BH',
         'SIS-LASSO-oracle-BH', 'SIS-SCAD-oracle-BH',
         'SIS-SLOPE', 'SIS-Knockoff',
         'db-ISIS-LASSO-bootWY', 'db-SIS-LASSO-BH', 'db-SIS-LASSO-bootWY',
         'SIS-LASSO', 'SIS-SCAD')
res <- resall %>% filter(!(Method %in% rmv)) 
res$Method <- fct_drop(res$Method)
unique(res$Method)

# check nubmer of obs
aggregate(nsel ~ ID + Method + Model + n + p + split + p.beta + beta0 + betaA + betaL + sigma + betaAsign + dsn + stdA + delta,
          data = res, FUN = length)

# pretty labels
res$nlab <- paste0('n = ', res$n)
res$plab <- factor(paste0('p = ', res$p), levels = unique(paste0('p = ', res$p)))
res$modlab <- ifelse(res$delta == 1, 'No Effect Modification', 'Strong Effect Modification')
res$split <- ifelse(res$split == 0.5, 0.6, res$split)
res$splitlab <- factor(paste0(res$split*100, '% Confounded'), 
                       levels = unique(paste0(res$split*100, '% Confounded')))
res$TN <- res$TNR * (res$p - res$p.beta)
res$ACC <- (res$TP + res$TN) / res$p
levels(res$Method) <- c('iterSIS-dbLASSO-BH',
                        'iterSIS-LASSO',
                        'iterSIS-SCAD',
                        'LASSO', 'SCAD')
res$model <- res$Model
levels(res$model) <- c('Select L', 'Select L EffMod', 
                       'Require L', 'Require L EffMod', 
                       'Conditional Std', 
                       'Ignore L', 'Ignore L EffMod')
res$model <- relevel(res$model, ref = 'Conditional Std')

# TPR plots
pdf('./figs/TPR_delta=-0.9_boxplot.pdf', width = 12)
ggplot(subset(res, delta == -0.9)) +
    geom_boxplot(aes(y = TPR, x = model, color = model)) +
    facet_grid(plab + splitlab ~ Method) +
    xlab('') + ylab('TPR') +
    scale_color_npg() +
    theme(axis.text.x = element_blank(),
          legend.position = 'bottom')
dev.off()
pdf('./figs/TPR_delta=1_boxplot.pdf', width = 12)
ggplot(subset(res, delta == 1)) +
    geom_boxplot(aes(y = TPR, x = model, color = model)) +
    facet_grid(plab + splitlab ~ Method) +
    xlab('') + ylab('TPR') +
    scale_color_npg() +
    theme(axis.text.x = element_blank(),
          legend.position = 'bottom')
dev.off()
pdf('./figs/TPR_delta=-0.9_violin.pdf', width = 12)
ggplot(subset(res, delta == -0.9)) +
    geom_violin(aes(y = TPR, x = model, color = model)) +
    facet_grid(plab + splitlab ~ Method) +
    xlab('') + ylab('TPR') +
    scale_color_npg() +
    theme(axis.text.x = element_blank(),
          legend.position = 'bottom')
dev.off()
pdf('./figs/TPR_delta=1_violin.pdf', width = 12)
ggplot(subset(res, delta == 1)) +
    geom_violin(aes(y = TPR, x = model, color = model)) +
    facet_grid(plab + splitlab ~ Method) +
    xlab('') + ylab('TPR') +
    scale_color_npg() +
    theme(axis.text.x = element_blank(),
          legend.position = 'bottom')
dev.off()

# FDP plots
pdf('./figs/FDP_delta=-0.9_boxplot.pdf', width = 12)
ggplot(subset(res, delta == -0.9)) +
    geom_boxplot(aes(y = FDP, x = model, color = model)) +
    facet_grid(plab + splitlab ~ Method) +
    xlab('') + ylab('FDP') +
    scale_color_npg() +
    theme(axis.text.x = element_blank(),
          legend.position = 'bottom')
dev.off()
pdf('./figs/FDP_delta=1_boxplot.pdf', width = 12)
ggplot(subset(res, delta == 1)) +
    geom_boxplot(aes(y = FDP, x = model, color = model)) +
    facet_grid(plab + splitlab ~ Method) +
    xlab('') + ylab('FDP') +
    scale_color_npg() +
    theme(axis.text.x = element_blank(),
          legend.position = 'bottom')
dev.off()
pdf('./figs/FDP_delta=-0.9_violin.pdf', width = 12)
ggplot(subset(res, delta == -0.9)) +
    geom_violin(aes(y = FDP, x = model, color = model)) +
    facet_grid(plab + splitlab ~ Method) +
    xlab('') + ylab('FDP') +
    scale_color_npg() +
    theme(axis.text.x = element_blank(),
          legend.position = 'bottom')
dev.off()
pdf('./figs/FDP_delta=1_violin.pdf', width = 12)
ggplot(subset(res, delta == 1)) +
    geom_violin(aes(y = FDP, x = model, color = model)) +
    facet_grid(plab + splitlab ~ Method) +
    xlab('') + ylab('FDP') +
    scale_color_npg() +
    theme(axis.text.x = element_blank(),
          legend.position = 'bottom')
dev.off()

# FPR plots
pdf('./figs/FPR_delta=-0.9_boxplot.pdf', width = 12)
ggplot(subset(res, delta == -0.9)) +
    geom_boxplot(aes(y = FPR, x = model, color = model)) +
    facet_grid(plab + splitlab ~ Method) +
    xlab('') + ylab('FPR') +
    scale_color_npg() +
    theme(axis.text.x = element_blank(),
          legend.position = 'bottom')
dev.off()
pdf('./figs/FPR_delta=1_boxplot.pdf', width = 12)
ggplot(subset(res, delta == 1)) +
    geom_boxplot(aes(y = FPR, x = model, color = model)) +
    facet_grid(plab + splitlab ~ Method) +
    xlab('') + ylab('FPR') +
    scale_color_npg() +
    theme(axis.text.x = element_blank(),
          legend.position = 'bottom')
dev.off()
pdf('./figs/FPR_delta=-0.9_violin.pdf', width = 12)
ggplot(subset(res, delta == -0.9)) +
    geom_violin(aes(y = FPR, x = model, color = model)) +
    facet_grid(plab + splitlab ~ Method) +
    xlab('') + ylab('FPR') +
    scale_color_npg() +
    theme(axis.text.x = element_blank(),
          legend.position = 'bottom')
dev.off()
pdf('./figs/FPR_delta=1_violin.pdf', width = 12)
ggplot(subset(res, delta == 1)) +
    geom_violin(aes(y = FPR, x = model, color = model)) +
    facet_grid(plab + splitlab ~ Method) +
    xlab('') + ylab('FPR') +
    scale_color_npg() +
    theme(axis.text.x = element_blank(),
          legend.position = 'bottom')
dev.off()

# Tables
# helper function for tables, output "mean (std err)"
stderr <- function(x) sd(x) / sqrt(length(x))
meanvar <- function(x, digits = 2, method = "SE") {
    m <- mean(x)
    rm <- sprintf(paste0("%.", digits, "f"), round(m, digits))
    if (method == "SE") den <- stderr(x) else den <- sd(x)
    rden <- sprintf(paste0("%.", digits, "f"), round(den, digits))
    out <- paste(rm," (", rden,")", sep="")
    return(out)
}
Mean <- function(x, digits = 2) sprintf(paste0("%.", digits, "f"), round(mean(x), digits))


# Make a FDP/TPR summary table
library(tables)
t1 <- tabular(Factor(modlab, 'Effect Modification') * Factor(plab, '\\# Features')  * Factor(splitlab, 'Relevant Features') * Factor(model, 'Model')  ~ 
                  Factor(Method) * (TPR+FPR) * Heading() * Mean,
              data = res, suppress = 1)
latex(t1, file = './tables/TPR-FPR.tex')

t2 <- tabular(Factor(modlab, 'Effect Modification') * Factor(plab, '\\# Features')  * Factor(splitlab, 'Relevant Features') * Factor(model, 'Model')  ~ 
                  Factor(Method) * (TPR+FDP) * Heading() * Mean,
              data = res, suppress = 1)
latex(t2, file = './tables/TPR-FDP.tex')

t3 <- tabular(Factor(modlab, 'Effect Modification') * Factor(plab, '\\# Features')  * Factor(splitlab, 'Relevant Features') * Factor(model, 'Model')  ~ 
                  Factor(Method) * TPR * Heading() * meanvar,
              data = res, suppress = 1)
latex(t3, file = './tables/TPR-se.tex')

t4 <- tabular(Factor(modlab, 'Effect Modification') * Factor(plab, '\\# Features')  * Factor(splitlab, 'Relevant Features') * Factor(model, 'Model')  ~ 
                  Factor(Method) * FDP * Heading() * meanvar,
              data = res, suppress = 1)
latex(t4, file = './tables/FDP-se.tex')


t5 <- tabular(Factor(modlab, 'Effect Modification') * Factor(plab, '\\# Features')  * Factor(splitlab, 'Relevant Features') * Factor(model, 'Model')  ~ 
                  Factor(Method) * FPR * Heading() * meanvar,
              data = res, suppress = 1)
latex(t5, file = './tables/FPR-se.tex')
```
