## Supplementary material for "Feature selection and causal analysis for microbiome studies in the presence of confounding using standardization": R code: 03_simulation_AUC.R

```
rm(list = ls())

set.seed(09202018)

require(SISmod)
require(hdi)
require(foreach)
require(doParallel)
devtools::session_info()
registerDoParallel(detectCores())


# Number of simulations
N <- 1:100
repIDs <- paste0('rep', N)
# Number of units (observations -- plots, plants, etc)
n <- c(50, 100)
# Number of OTUs/microbiome features
p <- c(50, 2000)
# Number of OTUs with an effect on response
p.beta <- 5
# Magntude, sign of p.beta, interactions?
betaA <- 3
betaAsign <- 'Mixed'
delta <- c(1, -0.9)
# Treatment effect
betaL <- 5
# Proportion of DA OTUs
prop.DA <- 0.3
# Split of p.beta amoung DA OTUs and nDA OTUs
split <- c(0, 0.5, 1)
# OTUs dsn
dsn <- c('Poisson', 'NB')
# Standardize X for data generation?
stdA <- TRUE
# Intercept
beta0 <- 3
# Variability (spherical)
sigma <- 0.25
conds <- expand.grid(n = n, p = p, split = split, p.beta = p.beta, 
                     beta0 = beta0, betaA = betaA, betaL = betaL, sigma = sigma,
                     betaAsign = betaAsign, dsn = dsn, stdA = stdA, delta = delta)
J <- nrow(conds)
conds$ID <- 1:J
J

# Functions to extract selected results.
sumdat <- function(res, p, std = FALSE) {
    nms <- names(res)
    ans <- lapply(1:length(res), function(r) {
        if (nms[r] %in% c('db-SIS-LASSO-BH', 'db-SIS-LASSO-BH', 'db-SIS-LASSO-bootWY',
                          'db-ISIS-LASSO-BH', 'db-ISIS-LASSO-BH', 'db-ISIS-LASSO-bootWY')) {
            if (class(res[[r]])[1] == 'logical') {
                ps <- pscorr <- betahat <- rep(NA, p)
            } else {
                if (std) {
                    bALL <- betahatALL <- rep(0, 2*p)
                    seALL <- rep(0, 2*p)
                    ids <- as.numeric(names(res[[r]]$pval))
                    bALL[ids] <- res[[r]]$bhat
                    betahatALL[ids] <- res[[r]]$betahat
                    seALL[ids] <- res[[r]]$se
                    b0 <- bALL[1:p]; b1 <- bALL[(p+1):(2*p)]
                    se0 <- seALL[1:p]; se1 <- seALL[(p+1):(2*p)]
                    betahat0 <- betahatALL[1:p]; betahat1 <- betahatALL[(p+1):(2*p)]
                    b <- (b0 + b1)
                    betahat <- (betahat0 + betahat1 ) / 2
                    se <- sqrt(se0^2 + se1^2)
                    z <- b / se
                    ps <- 2 * (1 - pnorm(abs(z)))
                    pscorr <- p.adjust(ps, method = 'fdr')
                } else {
                    ps <- pscorr <- rep(1, p)
                    betahat <- rep(0, p)
                    ids <- as.numeric(names(res[[r]]$pval))
                    ps[ids] <- res[[r]]$pval
                    pscorr[ids] <- res[[r]]$pval.corr
                    betahat[ids] <- res[[r]]$betahat
                }
            }
            df <- data.frame(pval = ps[1:p], pval.corr = pscorr[1:p], betahat = betahat[1:p],
                             OTU = 1:p, Method = nms[r])
            df$pval[is.nan(df$pval)] <- 1
            df$pval.corr[is.nan(df$pval.corr)] <- 1
        } else {
            df <- NULL
        }
        return(df)
    })
    ans <- do.call('rbind', ans)
    return(ans)
}
sumdatcond <- function(res0, res1, p) {
    nms <- names(res0)
    ans <- lapply(1:length(res0), function(r) {
        if (nms[r] %in% c('db-SIS-LASSO-BH', 'db-SIS-LASSO-BH', 'db-SIS-LASSO-bootWY',
                          'db-ISIS-LASSO-BH', 'db-ISIS-LASSO-BH', 'db-ISIS-LASSO-bootWY')) {
                b1 <- b0 <- rep(0, p)
                betahat1 <- betahat0 <- rep(0, p)
                se1 <- se0 <- rep(0, p)
                if (class(res0[[r]])[1] == 'logical') {
                    b0 <- betahat0 <- rep(NA, p)
                } else {
                    ids0 <- as.numeric(names(res0[[r]]$pval))
                    b0[ids0] <- res0[[r]]$bhat
                    betahat0[ids0] <- res0[[r]]$betahat
                    se0[ids0] <- res0[[r]]$se
                }
                if (class(res1[[r]])[1] == 'logical') {
                    b1 <- betahat1 <- rep(NA, p)
                } else {
                    ids1 <- as.numeric(names(res1[[r]]$pval))
                    b1[ids1] <- res1[[r]]$bhat
                    betahat1[ids1] <- res1[[r]]$betahat
                    se1[ids1] <- res1[[r]]$se
                }
                b <- (b0 + b1)
                betahat <- (betahat0 + betahat1 ) / 2
                se <- sqrt(se0^2 + se1^2)
                z <- b / se
                ps <- 2 * (1 - pnorm(abs(z)))
                ps[is.na(ps)] <- 1
                pscorr <- p.adjust(ps, method = 'fdr')
            df <- data.frame(pval = ps[1:p], pval.corr = pscorr[1:p], betahat = betahat[1:p],
                             OTU = 1:p, Method = nms[r])
        } else {
            df <- NULL
        }
        return(df)
    })
    ans <- do.call('rbind', ans)
    return(ans)
}

# Extract results for model selection.
o <- foreach(j = 1:J, .combine = rbind) %dopar% {
    load(paste0('./dump/data_params_ID', j, '.Rdat'))
    data <- out
    if (data[[repIDs[1]]]$params$ID != j) print('Condition ID does not match index')
    load(paste0('./dump/sim_fits_ID', j, '_dbLASSO.Rdat'))
    oj <- foreach(repID = repIDs, .combine = rbind) %do% {
        print(paste('Exracting p-values for condition ID', j, 'simulation', repID))
        thisft <- ft[[repID]]
        d <- data[[repID]]
        p <- d$params$p
        # Model 1: ignore treatment
        o1 <- sumdat(thisft$Model1, p = p, std = FALSE)
        o1$Model <- 'Ignore L'
        o1.int.std <- sumdat(thisft$Model1.int, p = p, std = TRUE)
        o1.int.std$Model <- 'Ignore L Std'
        # Model 2: additive treatment
        o2 <- sumdat(thisft$Model2, p = p, std = FALSE)
        o2$Model <- 'Additive L'
        o2.keepL <- sumdat(thisft$Model2.keepL, p = p, std = FALSE)
        o2.keepL$Model <- 'Additive Unpenalized L'
        o2.int.std <- sumdat(thisft$Model2.int, p = p, std = TRUE)
        o2.int.std$Model <- 'Additive L Std'
        o2.int.keepL.std <- sumdat(thisft$Model2.int.keepL, p = p, std = TRUE)
        o2.int.keepL.std$Model <- 'Additive Unpenalized L Std'
        # Model 3: conditional on treatment
        o3 <- sumdatcond(thisft$Model3.0, thisft$Model3.1, p = p)
        o3$Model <- 'Conditional Std'
        library(pROC)
        out <- list(Ignore = o1, Additive = o2, Additive.keepL = o2.keepL,
                    Ignore.int.std = o1.int.std, Additive.int.std = o2.int.std, 
                    Additive.int.keepL.std = o2.int.keepL.std, Conditional.std = o3)
        out <- do.call('rbind', out)
        out$ID <- d$params$ID
        out$nsim <- repID
        out$n <- d$params$n
        out$p <- d$params$p
        out$split <- d$params$split
        out$p.beta <- d$params$p.beta
        out$beta0 <- d$params$beta0
        out$betaA <- d$params$betaA
        out$betaL <- d$params$betaL
        out$sigma <- d$params$sigma
        out$betaAsign <- d$params$betaAsign
        out$dsn <- d$params$dsn
        out$stdA <- d$params$stdA
        out$delta <- d$params$delta
        out$truth <- as.numeric((d$beta[1:p] != 0))
        out
    }
    write.csv(oj, file = paste0('./dump/results_ID', j, '_dbLASSO.Rdat'), row.names = FALSE)
    NULL
}


# AUC

require(pROC)
require(dplyr)
require(ggplot2)
clrs <- c('#e41a1c','#377eb8','#a65628','#4daf4a','#984ea3','#ff7f00','#999999')
theme_set(theme_bw(base_size = 14))

get.auc <- function(true, pred, partial = FALSE) {
    if(any(is.na(pred))) return(NA)
    if (partial) {
        ans <- tryCatch(auc(roc(true, pred), partial.auc = c(1,0.9)),
                        error = function(e) NA)
    } else { 
        ans <- tryCatch(auc(roc(true, pred)),
                        error = function(e) NA)
    }
    return(as.numeric(ans))
}

o <- foreach(j = 1:J, .combine = rbind) %dopar% {
    oj <- read.csv(paste0('./dump/results_ID', j, '_dbLASSO.Rdat'))
    AUC <- oj %>% 
        group_by(Method, Model, nsim, ID, n, p, split, p.beta, beta0, betaA,
                 betaL, sigma, betaAsign, dsn, stdA, delta, ID) %>%
        summarise(auc = get.auc(truth, pval),
                  pauc = get.auc(truth, pval, TRUE))
    AUC
}


o$nlab <- paste0('n = ', o$n)
o$plab <- factor(paste0('p = ', o$p), levels = unique(paste0('p = ', o$p)))
o$modlab <- factor(ifelse(o$delta == 1, 'No Effect Modifier', 'Strong Effect Modifier'))
o$modlab <- relevel(o$modlab, ref = 'Strong Effect Modifier')
o$split <- ifelse(o$split == 0.5, 0.6, o$split)
o$splitlab <- factor(paste0(o$split*100, '% Confounded'), levels = unique(paste0(o$split*100, '% Confounded')))
o$model <- factor(o$Model, labels = c('Select L', 'Select L EffMod', 'Require L', 'Require L EffMod', 
                                      'Conditional Std', 'Ignore L', 'Ignore L EffMod'))
o$model <- relevel(o$model, ref = 'Conditional Std')


write.csv(o, file = './results_AUC_dbLASSO.csv', row.names = FALSE)
o <- read.csv(file = './results_AUC_dbLASSO.csv')

pdf('./figs/AUC_box_n=50_Pois_flip.pdf', width = 8, height = 12)
ggplot(subset(o,  n == 50 & dsn == 'Poisson')) +
    geom_boxplot(aes(y = auc, x = model, colour = model, fill = model)) +
    facet_grid( modlab + plab ~ splitlab) +
    xlab('Model') + ylab('AUC') +
    theme(legend.position = 'none') +
    scale_colour_manual(values = clrs) +
    scale_fill_manual(values = alpha(clrs, 0.25)) +
    theme(axis.text.x = element_text(angle = 60, hjust = 1))
dev.off()


pdf('./figs/AUC_box_n=50_Pois.pdf', width = 8)
ggplot(subset(o,  n == 50 & dsn == 'Poisson')) +
    geom_boxplot(aes(y = auc, x = model, colour = model, fill = model)) +
    facet_grid(splitlab ~ modlab + plab) +
    xlab('Model') + ylab('AUC') +
    theme(legend.position = 'none') +
    scale_colour_manual(values = clrs) +
    scale_fill_manual(values = alpha(clrs, 0.25)) +
    theme(axis.text.x = element_text(angle = 60, hjust = 1))
dev.off()


pdf('./figs/AUC_box_n=100_Pois_flip.pdf', width = 8, height = 12)
ggplot(subset(o,  n == 100 & dsn == 'Poisson')) +
    geom_boxplot(aes(y = auc, x = model, colour = model, fill = model)) +
    facet_grid(modlab + plab ~ splitlab) +
    xlab('Model') + ylab('AUC') +
    theme(legend.position = 'none') +
    scale_colour_manual(values = clrs) +
    scale_fill_manual(values = alpha(clrs, 0.25)) +
    theme(axis.text.x = element_text(angle = 60, hjust = 1))
dev.off()


pdf('./figs/AUC_box_n=100_Pois.pdf', width = 8)
ggplot(subset(o,  n == 100 & dsn == 'Poisson')) +
    geom_boxplot(aes(y = auc, x = model, colour = model, fill = model)) +
    facet_grid(splitlab ~ modlab + plab) +
    xlab('Model') + ylab('AUC') +
    theme(legend.position = 'none') +
    scale_colour_manual(values = clrs) +
    scale_fill_manual(values = alpha(clrs, 0.25)) +
    theme(axis.text.x = element_text(angle = 60, hjust = 1))
dev.off()

pdf('./figs/AUC_box_n=50_NB_flip.pdf', width = 8, height = 12)
ggplot(subset(o,  n == 50 & dsn == 'NB')) +
    geom_boxplot(aes(y = auc, x = model, colour = model, fill = model)) +
    facet_grid(modlab + plab ~ splitlab) +
    xlab('Model') + ylab('AUC') +
    theme(legend.position = 'none') +
    scale_colour_manual(values = clrs) +
    scale_fill_manual(values = alpha(clrs, 0.25)) +
    theme(axis.text.x = element_text(angle = 60, hjust = 1))
dev.off()

pdf('./figs/AUC_box_n=50_NB.pdf', width = 8)
ggplot(subset(o,  n == 50 & dsn == 'NB')) +
    geom_boxplot(aes(y = auc, x = model, colour = model, fill = model)) +
    facet_grid(splitlab ~ modlab + plab) +
    xlab('Model') + ylab('AUC') +
    theme(legend.position = 'none') +
    scale_colour_manual(values = clrs) +
    scale_fill_manual(values = alpha(clrs, 0.25)) +
    theme(axis.text.x = element_text(angle = 60, hjust = 1))
dev.off()


pdf('./figs/AUC_box_n=100_NB_flip.pdf', width = 8, height = 12)
ggplot(subset(o,  n == 100 & dsn == 'NB')) +
    geom_boxplot(aes(y = auc, x = model, colour = model, fill = model)) +
    facet_grid(modlab + plab ~ splitlab) +
    xlab('Model') + ylab('AUC') +
    theme(legend.position = 'none') +
    scale_colour_manual(values = clrs) +
    scale_fill_manual(values = alpha(clrs, 0.25)) +
    theme(axis.text.x = element_text(angle = 60, hjust = 1))
dev.off()


pdf('./figs/AUC_box_n=100_NB.pdf', width = 8)
ggplot(subset(o,  n == 100 & dsn == 'NB')) +
    geom_boxplot(aes(y = auc, x = model, colour = model, fill = model)) +
    facet_grid(splitlab ~ modlab + plab) +
    xlab('Model') + ylab('AUC') +
    theme(legend.position = 'none') +
    scale_colour_manual(values = clrs) +
    scale_fill_manual(values = alpha(clrs, 0.25)) +
    theme(axis.text.x = element_text(angle = 60, hjust = 1))
dev.off()

pdf('./figs/AUC_box_NB.pdf', width = 12)
ggplot(subset(o,  dsn == 'NB')) +
    geom_boxplot(aes(y = auc, x = model, colour = model, fill = model)) +
    facet_grid(nalb + splitlab ~ modlab + plab) +
    xlab('Model') + ylab('AUC') +
    theme(legend.position = 'none') +
    scale_colour_manual(values = clrs) +
    scale_fill_manual(values = alpha(clrs, 0.25)) +
    theme(axis.text.x = element_text(angle = 60, hjust = 1))
dev.off()

pdf('./figs/AUC_box_Pois.pdf', width = 12)
ggplot(subset(o,  dsn == 'Poisson')) +
    geom_boxplot(aes(y = auc, x = model, colour = model, fill = model)) +
    facet_grid(nlab + splitlab ~ modlab + plab) +
    xlab('Model') + ylab('AUC') +
    theme(legend.position = 'none') +
    scale_colour_manual(values = clrs) +
    scale_fill_manual(values = alpha(clrs, 0.25)) +
    theme(axis.text.x = element_text(angle = 60, hjust = 1))
dev.off()


# error rate control
alpha <- seq(0, 0.1, by = 0.01)
fp <- function(truth, pval, alpha, p.adjust = 'none') {
    pval[is.na(pval)] <- 1
    pcor <- p.adjust(pval, p.adjust)
    rejectH0 <- (pcor < alpha)
    return(sum(truth[rejectH0] == 0))
}
fdp <- function(truth, pval, alpha, p.adjust = 'none') {
    pval[is.na(pval)] <- 1
    pcor <- p.adjust(pval, p.adjust)
    rejectH0 <- (pcor < alpha)
    if (sum(rejectH0) == 0) {
        out <- 0
    } else {
        npos <- sum(rejectH0)
        fpos <- sum(truth[rejectH0] == 0)
        out <- fpos / npos
    }
    return(out)
}
e <- foreach(j = 1:J, .combine = rbind) %dopar% {
    oj <- read.csv(paste0('./dump/results_ID', j, '_dbLASSO.Rdat'))
    out <- foreach(a = alpha, .combine = rbind) %do% {
        outa <- oj %>% 
            group_by(Method, Model, nsim, ID, n, p, split, p.beta, beta0, betaA,
                     betaL, sigma, betaAsign, dsn, stdA, delta) %>%
            summarise(FP.BH = fp(truth, pval, a, 'BH'),
                      FP.BY = fp(truth, pval, a, 'BY'),
                      FDP.BH = fdp(truth, pval, a, 'BH'),
                      FDP.BY = fdp(truth, pval, a, 'BY'))
        outa$alpha <- a
        outa
    }
    out
}

e$nlab <- paste0('n = ', e$n)
e$plab <- factor(paste0('p = ', e$p), levels = unique(paste0('p = ', e$p)))
e$modlab <- factor(ifelse(e$delta == 1, 'No Effect Modifier', 'Strong Effect Modifier'))
e$modlab <- relevel(e$modlab, ref = 'Strong Effect Modifier')
e$split <- ifelse(e$split == 0.5, 0.6, e$split)
e$splitlab <- factor(paste0(e$split*100, '% Confounded'), levels = unique(paste0(e$split*100, '% Confounded')))
e$model <- factor(e$Model, labels = c('Select L', 'Select L EffMod', 'Require L', 'Require L EffMod', 
                                      'Conditional Std', 'Ignore L', 'Ignore L EffMod'))
e$model <- relevel(e$model, ref = 'Conditional Std')
e$FP.BH[is.na(e$FP.BH)] <- 0
e$FP.BY[is.na(e$FP.BY)] <- 0
e$anyFP.BH <- as.numeric(e$FP.BH >= 1)
e$anyFP.BY <- as.numeric(e$FP.BY >= 1)


write.csv(e, file = './results_FDR_dbLASSO.csv', row.names = FALSE)
#e <- read.csv(file = './results_FDR_dbLASSO.csv')


pdf('./figs/FDP-BH_n=50_Pois.pdf_flip.pdf', width = 8, height = 12)
ggplot(subset(e, n == 50 & dsn == 'Poisson')) +
    stat_summary(aes(y = FDP.BH, x = alpha, color = model), 
                 fun.y = 'mean', geom = 'line') +
    geom_abline(intercept = 0, slope = 1) +
    facet_grid(modlab + plab ~ splitlab, scales = "free_y") +
    xlab('Nominal FDR') + ylab('Mean FDP') +
    theme(legend.position = 'bottom') +
    guides(colour = guide_legend(nrow = 1)) +
    scale_x_continuous(breaks = c(0, 0.025, 0.05, 0.075, 0.1), 
                       labels = c(0, 0.025, 0.05, 0.075, 0.1)) +
    labs(color = 'Model') +
    scale_colour_manual(values = clrs) +
    guides(colour=guide_legend(nrow=2,byrow=TRUE))
dev.off()


pdf('./figs/FDP-BH_n=50_Pois.pdf', width = 12)
ggplot(subset(e, n == 50 & dsn == 'Poisson')) +
    stat_summary(aes(y = FDP.BH, x = alpha, color = model), 
                 fun.y = 'mean', geom = 'line') +
    geom_abline(intercept = 0, slope = 1) +
    facet_grid(splitlab ~ modlab + plab) +
    xlab('Nominal FDR') + ylab('Mean FDP') +
    theme(legend.position = 'bottom') +
    guides(colour = guide_legend(nrow = 1)) +
    scale_x_continuous(breaks = c(0, 0.025, 0.05, 0.075, 0.1), 
                       labels = c(0, 0.025, 0.05, 0.075, 0.1)) +
    labs(color = 'Model') +
        scale_colour_manual(values = clrs)
dev.off()


pdf('./figs/FDP-BH_n=100_Pois_flip.pdf', width = 8, height = 12)
ggplot(subset(e, n == 100 & dsn == 'Poisson')) +
    stat_summary(aes(y = FDP.BH, x = alpha, color = model), 
                 fun.y = 'mean', geom = 'line') +
    geom_abline(intercept = 0, slope = 1) +
    facet_grid(modlab + plab ~ splitlab, scales = "free_y") +
    xlab('Nominal FDR') + ylab('Mean FDP') +
    theme(legend.position = 'bottom') +
    guides(colour = guide_legend(nrow = 1)) +
    scale_x_continuous(breaks = c(0, 0.025, 0.05, 0.075, 0.1), 
                       labels = c(0, 0.025, 0.05, 0.075, 0.1)) +
    labs(color = 'Model') +
    scale_colour_manual(values = clrs) +
    guides(colour=guide_legend(nrow=2,byrow=TRUE))
dev.off()


pdf('./figs/FDP-BH_n=100_Pois.pdf', width = 12)
ggplot(subset(e, n == 100 & dsn == 'Poisson')) +
    stat_summary(aes(y = FDP.BH, x = alpha, color = model), 
                 fun.y = 'mean', geom = 'line') +
    geom_abline(intercept = 0, slope = 1) +
    facet_grid(splitlab ~ modlab + plab) +
    xlab('Nominal FDR') + ylab('Mean FDP') +
    theme(legend.position = 'bottom') +
    guides(colour = guide_legend(nrow = 1)) +
    scale_x_continuous(breaks = c(0, 0.025, 0.05, 0.075, 0.1), 
                       labels = c(0, 0.025, 0.05, 0.075, 0.1)) +
    labs(color = 'Model') +
        scale_colour_manual(values = clrs)
dev.off()


pdf('./figs/FDP-BH_n=50_NB_flip.pdf', width = 8, height = 12)
ggplot(subset(e, n == 50 & dsn == 'NB')) +
    stat_summary(aes(y = FDP.BH, x = alpha, color = model), 
                 fun.y = 'mean', geom = 'line') +
    geom_abline(intercept = 0, slope = 1) +
    facet_grid(modlab + plab ~ splitlab, scales = "free_y") +
    xlab('Nominal FDR') + ylab('Mean FDP') +
    theme(legend.position = 'bottom') +
    guides(colour = guide_legend(nrow = 1)) +
    scale_x_continuous(breaks = c(0, 0.025, 0.05, 0.075, 0.1), 
                       labels = c(0, 0.025, 0.05, 0.075, 0.1)) +
    labs(color = 'Model') +
    scale_colour_manual(values = clrs) +
    guides(colour=guide_legend(nrow=2,byrow=TRUE))
dev.off()


pdf('./figs/FDP-BH_n=50_NB.pdf', width = 12)
ggplot(subset(e, n == 50 & dsn == 'NB')) +
    stat_summary(aes(y = FDP.BH, x = alpha, color = model), 
                 fun.y = 'mean', geom = 'line') +
    geom_abline(intercept = 0, slope = 1) +
    facet_grid(splitlab ~ modlab + plab) +
    xlab('Nominal FDR') + ylab('Mean FDP') +
    theme(legend.position = 'bottom') +
    guides(colour = guide_legend(nrow = 1)) +
    scale_x_continuous(breaks = c(0, 0.025, 0.05, 0.075, 0.1),
                       labels = c(0, 0.025, 0.05, 0.075, 0.1)) +
    labs(color = 'Model') +
        scale_colour_manual(values = clrs)
dev.off()


pdf('./figs/FDP-BH_n=100_NB_flip.pdf', width = 8, height = 12)
ggplot(subset(e, n == 100 & dsn == 'NB')) +
    stat_summary(aes(y = FDP.BH, x = alpha, color = model), 
                 fun.y = 'mean', geom = 'line') +
    geom_abline(intercept = 0, slope = 1) +
    facet_grid(modlab + plab ~ splitlab, scales = "free_y") +
    xlab('Nominal FDR') + ylab('Mean FDP') +
    theme(legend.position = 'bottom') +
    guides(colour = guide_legend(nrow = 1)) +
    scale_x_continuous(breaks = c(0, 0.025, 0.05, 0.075, 0.1), 
                       labels = c(0, 0.025, 0.05, 0.075, 0.1)) +
    labs(color = 'Model') +
    scale_colour_manual(values = clrs) +
    guides(colour=guide_legend(nrow=2,byrow=TRUE))
dev.off()


pdf('./figs/FDP-BH_n=100_NB.pdf', width = 12)
ggplot(subset(e, n == 100 & dsn == 'NB')) +
    stat_summary(aes(y = FDP.BH, x = alpha, color = model), 
                 fun.y = 'mean', geom = 'line') +
    geom_abline(intercept = 0, slope = 1) +
    facet_grid(splitlab ~ modlab + plab) +
    xlab('Nominal FDR') + ylab('Mean FDP') +
    theme(legend.position = 'bottom') +
    guides(colour = guide_legend(nrow = 1)) +
    scale_x_continuous(breaks = c(0, 0.025, 0.05, 0.075, 0.1),
                       labels = c(0, 0.025, 0.05, 0.075, 0.1)) +
    labs(color = 'Model') +
        scale_colour_manual(values = clrs)
dev.off()


pdf('./figs/FDP-BH_NB.pdf', width = 12)
ggplot(subset(e, dsn == 'NB')) +
    stat_summary(aes(y = FDP.BH, x = alpha, color = model), 
                 fun.y = 'mean', geom = 'line') +
    geom_abline(intercept = 0, slope = 1) +
    facet_grid(nlab + splitlab ~ modlab + plab) +
    xlab('Nominal FDR') + ylab('Mean FDP') +
    theme(legend.position = 'bottom') +
    guides(colour = guide_legend(nrow = 1)) +
    scale_x_continuous(breaks = c(0, 0.025, 0.05, 0.075, 0.1),
                       labels = c(0, 0.025, 0.05, 0.075, 0.1)) +
    labs(color = 'Model') +
        scale_colour_manual(values = clrs)
dev.off()


pdf('./figs/FDP-BH_Pois.pdf', width = 12)
ggplot(subset(e, dsn == 'Poisson')) +
    stat_summary(aes(y = FDP.BH, x = alpha, color = model),
                 fun.y = 'mean', geom = 'line') +
    geom_abline(intercept = 0, slope = 1) +
    facet_grid(nlab + splitlab ~ modlab + plab) +
    xlab('Nominal FDR') + ylab('Mean FDP') +
    theme(legend.position = 'bottom') +
    guides(colour = guide_legend(nrow = 1)) +
    scale_x_continuous(breaks = c(0, 0.025, 0.05, 0.075, 0.1), 
                       labels = c(0, 0.025, 0.05, 0.075, 0.1)) +
    labs(color = 'Model') +
    scale_colour_manual(values = clrs)
dev.off()


pdf('./figs/FDP-BY_n=50_Pois_flip.pdf', width = 8, height = 12)
ggplot(subset(e, n == 50 & dsn == 'Poisson')) +
    stat_summary(aes(y = FDP.BY, x = alpha, color = model),
                 fun.y = 'mean', geom = 'line') +
    geom_abline(intercept = 0, slope = 1) +
    facet_grid(modlab + plab ~ splitlab, scales = "free_y") +
    xlab('Nominal FDR') + ylab('Mean FDP') +
    theme(legend.position = 'bottom') +
    guides(colour = guide_legend(nrow = 1)) +
    scale_x_continuous(breaks = c(0, 0.025, 0.05, 0.075, 0.1), 
                       labels = c(0, 0.025, 0.05, 0.075, 0.1)) +
    labs(color = 'Model') +
    scale_colour_manual(values = clrs) +
    guides(colour=guide_legend(nrow=2,byrow=TRUE))
dev.off()

pdf('./figs/FDP-BY_n=50_Pois.pdf', width = 12)
ggplot(subset(e, n == 50 & dsn == 'Poisson')) +
    stat_summary(aes(y = FDP.BY, x = alpha, color = model),
                 fun.y = 'mean', geom = 'line') +
    geom_abline(intercept = 0, slope = 1) +
    facet_grid(splitlab ~ modlab + plab) +
    xlab('Nominal FDR') + ylab('Mean FDP') +
    theme(legend.position = 'bottom') +
    guides(colour = guide_legend(nrow = 1)) +
    scale_x_continuous(breaks = c(0, 0.025, 0.05, 0.075, 0.1), 
                       labels = c(0, 0.025, 0.05, 0.075, 0.1)) +
    labs(color = 'Model') +
    scale_colour_manual(values = clrs)
dev.off()


pdf('./figs/FDP-BY_n=100_Pois_flip.pdf', width = 8, height = 12)
ggplot(subset(e, n == 100 & dsn == 'Poisson')) +
    stat_summary(aes(y = FDP.BY, x = alpha, color = model),
                 fun.y = 'mean', geom = 'line') +
    geom_abline(intercept = 0, slope = 1) +
    facet_grid(modlab + plab ~ splitlab, scales = "free_y") +
    xlab('Nominal FDR') + ylab('Mean FDP') +
    theme(legend.position = 'bottom') +
    guides(colour = guide_legend(nrow = 1)) +
    scale_x_continuous(breaks = c(0, 0.025, 0.05, 0.075, 0.1), 
                       labels = c(0, 0.025, 0.05, 0.075, 0.1)) +
    labs(color = 'Model') +
    scale_colour_manual(values = clrs) +
    guides(colour=guide_legend(nrow=2,byrow=TRUE))
dev.off()


pdf('./figs/FDP-BY_n=100_Pois.pdf', width = 12)
ggplot(subset(e, n == 100 & dsn == 'Poisson')) +
    stat_summary(aes(y = FDP.BY, x = alpha, color = model),
                 fun.y = 'mean', geom = 'line') +
    geom_abline(intercept = 0, slope = 1) +
    facet_grid(splitlab ~ modlab + plab) +
    xlab('Nominal FDR') + ylab('Mean FDP') +
    theme(legend.position = 'bottom') +
    guides(colour = guide_legend(nrow = 1)) +
    scale_x_continuous(breaks = c(0, 0.025, 0.05, 0.075, 0.1), 
                       labels = c(0, 0.025, 0.05, 0.075, 0.1)) +
    labs(color = 'Model') +
    scale_colour_manual(values = clrs)
dev.off()


pdf('./figs/FDP-BY_n=50_NB_flip.pdf', width = 8, height = 12)
ggplot(subset(e, n == 50 & dsn == 'NB')) +
    stat_summary(aes(y = FDP.BY, x = alpha, color = model),
                 fun.y = 'mean', geom = 'line') +
    geom_abline(intercept = 0, slope = 1) +
    facet_grid(modlab + plab ~ splitlab, scales = "free_y") +
    xlab('Nominal FDR') + ylab('Mean FDP') +
    theme(legend.position = 'bottom') +
    guides(colour = guide_legend(nrow = 1)) +
    scale_x_continuous(breaks = c(0, 0.025, 0.05, 0.075, 0.1), 
                       labels = c(0, 0.025, 0.05, 0.075, 0.1)) +
    labs(color = 'Model') +
    scale_colour_manual(values = clrs) +
    guides(colour=guide_legend(nrow=2,byrow=TRUE))
dev.off()


pdf('./figs/FDP-BY_n=50_NB.pdf', width = 12)
ggplot(subset(e, n == 50 & dsn == 'NB')) +
    stat_summary(aes(y = FDP.BY, x = alpha, color = model), 
                 fun.y = 'mean', geom = 'line') +
    geom_abline(intercept = 0, slope = 1) +
    facet_grid(splitlab ~ modlab + plab) +
    xlab('Nominal FDR') + ylab('Mean FDP') +
    theme(legend.position = 'bottom') +
    guides(colour = guide_legend(nrow = 1)) +
    scale_x_continuous(breaks = c(0, 0.025, 0.05, 0.075, 0.1), 
                       labels = c(0, 0.025, 0.05, 0.075, 0.1)) +
    labs(color = 'Model') +
    scale_colour_manual(values = clrs)
dev.off()


pdf('./figs/FDP-BY_n=100_NB_flip.pdf', width = 8, height = 12)
ggplot(subset(e, n == 100 & dsn == 'NB')) +
    stat_summary(aes(y = FDP.BY, x = alpha, color = model),
                 fun.y = 'mean', geom = 'line') +
    geom_abline(intercept = 0, slope = 1) +
    facet_grid(modlab + plab ~ splitlab, scales = "free_y") +
    xlab('Nominal FDR') + ylab('Mean FDP') +
    theme(legend.position = 'bottom') +
    guides(colour = guide_legend(nrow = 1)) +
    scale_x_continuous(breaks = c(0, 0.025, 0.05, 0.075, 0.1), 
                       labels = c(0, 0.025, 0.05, 0.075, 0.1)) +
    labs(color = 'Model') +
    scale_colour_manual(values = clrs) +
    guides(colour=guide_legend(nrow=2,byrow=TRUE))
dev.off()


pdf('./figs/FDP-BY_n=100_NB.pdf', width = 12)
ggplot(subset(e, n == 100 & dsn == 'NB')) +
    stat_summary(aes(y = FDP.BY, x = alpha, color = model), 
                 fun.y = 'mean', geom = 'line') +
    geom_abline(intercept = 0, slope = 1) +
    facet_grid(splitlab ~ modlab + plab) +
    xlab('Nominal FDR') + ylab('Mean FDP') +
    theme(legend.position = 'bottom') +
    guides(colour = guide_legend(nrow = 1)) +
    scale_x_continuous(breaks = c(0, 0.025, 0.05, 0.075, 0.1),
                       labels = c(0, 0.025, 0.05, 0.075, 0.1)) +
    labs(color = 'Model') +
    scale_colour_manual(values = clrs)
dev.off()


pdf('./figs/FDP-BY_NB.pdf', width = 12)
ggplot(subset(e, dsn == 'NB')) +
    stat_summary(aes(y = FDP.BY, x = alpha, color = model),
                 fun.y = 'mean', geom = 'line') +
    geom_abline(intercept = 0, slope = 1) +
    facet_grid(nlab + splitlab ~ modlab + plab) +
    xlab('Nominal FDR') + ylab('Mean FDP') +
    theme(legend.position = 'bottom') +
    guides(colour = guide_legend(nrow = 1)) +
    scale_x_continuous(breaks = c(0, 0.025, 0.05, 0.075, 0.1), 
                       labels = c(0, 0.025, 0.05, 0.075, 0.1)) +
    labs(color = 'Model') +
    scale_colour_manual(values = clrs)
dev.off()


pdf('./figs/FDP-BY_Pois.pdf', width = 12)
ggplot(subset(e, dsn == 'Poisson')) +
    stat_summary(aes(y = FDP.BY, x = alpha, color = model), 
                 fun.y = 'mean', geom = 'line') +
    geom_abline(intercept = 0, slope = 1) +
    facet_grid(nlab + splitlab ~ modlab + plab) +
    xlab('Nominal FDR') + ylab('Mean FDP') +
    theme(legend.position = 'bottom') +
    guides(colour = guide_legend(nrow = 1)) +
    scale_x_continuous(breaks = c(0, 0.025, 0.05, 0.075, 0.1), 
                       labels = c(0, 0.025, 0.05, 0.075, 0.1)) +
    labs(color = 'Model') +
    scale_colour_manual(values = clrs)
dev.off()


##### addition 20190922 per HH's advice to look at delta from nomial

e_sum <- e %>%
    group_by(Method, Model, ID, n, p, split, p.beta, 
             beta0, betaA, betaL, sigma, betaAsign,
             dsn, stdA, delta, alpha, nlab, plab, modlab,
             splitlab, model) %>%
    summarise(
        mean.FDP.BH = mean(FDP.BH),
        mean.FDP.BY = mean(FDP.BY),
        mean.FDP.BH.MPE = mean((FDP.BH - alpha)/alpha),
        mean.FDP.BY.MPE = mean((FDP.BY - alpha)/alpha))
e_sum <- subset(e_sum, alpha >= 0.02)

pdf('./figs/FDP-MPE-BY_n=50_Pois.pdf', width = 12)
ggplot(subset(e_sum, n == 50 & dsn == 'Poisson')) +
    geom_line(aes(y = mean.FDP.BY.MPE, x = alpha, color = model), size = 1) +
    geom_hline(yintercept = 0, color = "black", size = 1) +
    facet_grid(splitlab ~ modlab + plab) +
    xlab('Nominal FDR') + ylab('FDP: mean proportion deviation from nomial') +
    theme(legend.position = 'bottom') +
    guides(colour = guide_legend(nrow = 1)) +
    scale_x_continuous(breaks = c(0, 0.025, 0.05, 0.075, 0.1), 
                       labels = c(0, 0.025, 0.05, 0.075, 0.1)) +
    labs(color = 'Model') +
    scale_colour_manual(values = clrs)
dev.off()

pdf('./figs/FDP-MPE-BY_n=100_Pois.pdf', width = 12)
ggplot(subset(e_sum, n == 100 & dsn == 'Poisson')) +
    geom_line(aes(y = mean.FDP.BY.MPE, x = alpha, color = model), size = 1) +
    geom_hline(yintercept = 0, color = "black", size = 1) +
    facet_grid(splitlab ~ modlab + plab) +
    xlab('Nominal FDR') + ylab('FDP: mean proportion deviation from nomial') +
    theme(legend.position = 'bottom') +
    guides(colour = guide_legend(nrow = 1)) +
    scale_x_continuous(breaks = c(0, 0.025, 0.05, 0.075, 0.1), 
                       labels = c(0, 0.025, 0.05, 0.075, 0.1)) +
    labs(color = 'Model') +
    scale_colour_manual(values = clrs)
dev.off()


pdf('./figs/FDP-MPE-BY_n=50_NB.pdf', width = 12)
ggplot(subset(e_sum, n == 50 & dsn == 'NB')) +
    geom_line(aes(y = mean.FDP.BY.MPE, x = alpha, color = model), size = 1) +
    geom_hline(yintercept = 0, color = "black", size = 1) +
    facet_grid(splitlab ~ modlab + plab) +
    xlab('Nominal FDR') + ylab('FDP: mean proportion deviation from nomial') +
    theme(legend.position = 'bottom') +
    guides(colour = guide_legend(nrow = 1)) +
    scale_x_continuous(breaks = c(0, 0.025, 0.05, 0.075, 0.1), 
                       labels = c(0, 0.025, 0.05, 0.075, 0.1)) +
    labs(color = 'Model') +
    scale_colour_manual(values = clrs)
dev.off()

pdf('./figs/FDP-MPE-BY_n=100_NB.pdf', width = 12)
ggplot(subset(e_sum, n == 100 & dsn == 'NB')) +
    geom_line(aes(y = mean.FDP.BY.MPE, x = alpha, color = model), size = 1) +
    geom_hline(yintercept = 0, color = "black", size = 1) +
    facet_grid(splitlab ~ modlab + plab) +
    xlab('Nominal FDR') + ylab('FDP: mean proportion deviation from nomial') +
    theme(legend.position = 'bottom') +
    guides(colour = guide_legend(nrow = 1)) +
    scale_x_continuous(breaks = c(0, 0.025, 0.05, 0.075, 0.1), 
                       labels = c(0, 0.025, 0.05, 0.075, 0.1)) +
    labs(color = 'Model') +
    scale_colour_manual(values = clrs)
dev.off()


pdf('./figs/FDP-MPE-BY_Pois.pdf', width = 12)
ggplot(subset(e_sum, dsn == 'NB')) +
    geom_line(aes(y = mean.FDP.BY.MPE, x = alpha, color = model), size = 1) +
    geom_hline(yintercept = 0, color = "black", size = 1) +
    facet_grid(nlab + splitlab ~ modlab + plab) +
    xlab('Nominal FDR') + ylab('FDP: mean proportion deviation from nomial') +
    theme(legend.position = 'bottom') +
    guides(colour = guide_legend(nrow = 1)) +
    scale_x_continuous(breaks = c(0, 0.025, 0.05, 0.075, 0.1), 
                       labels = c(0, 0.025, 0.05, 0.075, 0.1)) +
    labs(color = 'Model') +
    scale_colour_manual(values = clrs)
dev.off()


pdf('./figs/FDP-MPE-BY_NB.pdf', width = 12)
ggplot(subset(e_sum, dsn == 'Poisson')) +
    geom_line(aes(y = mean.FDP.BY.MPE, x = alpha, color = model), size = 1) +
    geom_hline(yintercept = 0, color = "black", size = 1) +
    facet_grid(nlab + splitlab ~ modlab + plab) +
    xlab('Nominal FDR') + ylab('FDP: mean proportion deviation from nomial') +
    theme(legend.position = 'bottom') +
    guides(colour = guide_legend(nrow = 1)) +
    scale_x_continuous(breaks = c(0, 0.025, 0.05, 0.075, 0.1), 
                       labels = c(0, 0.025, 0.05, 0.075, 0.1)) +
    labs(color = 'Model') +
    scale_colour_manual(values = clrs)
dev.off()
```
