## Supplementary material for "Feature selection and causal analysis for microbiome studies in the presence of confounding using standardization": R code: 04_real_data_analysis.Rmd

````
---
title: "DOE 2015 Sorghum Microbiome Data - Nitrogen"
author: "Emily Goren"
date: "10/5/2018 updated 1/15/2019, 12/8/2019"
output: pdf_document
---

```{r setup, include=FALSE}
knitr::opts_chunk$set(echo = FALSE, cache = TRUE, 
                      dev = c('png', 'pdf'), fig.height = 10, fig.width = 12)
require(tidyverse)
require(edgeR)
require(knitr)
theme_set(theme_bw(base_size = 16))
```

```{r data, echo=FALSE}
dir <- REDACTED
## Note - code to read in and clean data is modified from CY

## loading metabolites data 
## row: sample, columns: metabolities, size: 67 * 9 
mb_dat <- readxl::read_xlsx(paste0(dir, "doe_metabo_phytoh_2015.xlsx"), sheet = 1)

## loading OTU data: 5585 * 81, OTU: 5584  
otu_dat <- readxl::read_xlsx(paste0(dir, "doe_mic_phytoh_2015.xlsx"), sheet  = 1)

## loading sample matching key 
micro_key <- readxl::read_xlsx(paste0(dir, "micro_key.xlsx"), sheet = 1)
## micro_key_2 is used for matching mb_dat sample 
micro_key_2 <- readxl::read_xlsx(paste0(dir, "micro_key.xlsx"), sheet = 2, col_names = FALSE)
names(micro_key_2) <- c("sample_name", "sample_name_metabolites", "plant", "genotype", "collection_year", "collection_month",
                        "collection_day", "sample_isolated_from", "rep", "collection_site_or_growth_conditions")


metabolites_samplesClean <- function() {
  ## use mb_dat sample ID and micro_key to identify the sample, & experiment 
  
  ## check if any duplicated samples ID in mb_dat 
  if (any(duplicated(mb_dat$Sample))) {
    cat("There are duplicated samples in metabolites data set: ", mb_dat$Sample[duplicated(mb_dat$Sample)])
  }
  
  ## replace all dot in micro_key to match the format in metabolites data set 
  micro_key_2$sample_name_metabolites <- as.vector(sapply(micro_key_2$sample_name_metabolites, function(s){gsub("\\.", "\\_", s)}))
  
  ## check if all metabolites sample name have corresponding matching key 
  if (!all(mb_dat$Sample %in% micro_key_2$sample_name_metabolites)) stop("Not all metabolites samples have matching key\n") 
  
  ## extract sample information 
  mb_dat_sample_info <- micro_key_2 %>% filter(sample_name_metabolites %in% mb_dat$Sample) %>% data.frame() 
  #save(mb_dat_sample_info, file = "./RData/mb_dat_sample_info.RData")
  return(mb_dat_sample_info)
}  


mb_dat_sample_info <- metabolites_samplesClean()

## OTU
## OTU data clean 
out_datClean <- function() {
  ## check duplicated sample names in otu_dat 
  if (any(duplicated(names(otu_dat)))) cat("There is duplicated samples in otu_dat.\n")
  ## Only extract the samples that have metabolites values 
  ## get the sample name in mb_dat_sample_info match the format in otu data 
  samples_in_metabolites <- gsub("SAMPLE_", "", mb_dat_sample_info$sample_name)
  otu_samples <- names(otu_dat) 
  otu_selected_samples_cols <- otu_samples[otu_samples %in% samples_in_metabolites]
  ## add first column: NAME 
  otu_selected_samples_cols_1 <- c(names(otu_dat)[1], otu_selected_samples_cols)
  ## selected OTUs 5585 * 68
  otu_selected <- otu_dat[, otu_selected_samples_cols_1]
  ## remove first row : class 
  otu_selected <- otu_selected[-1, ]
  ## convert all to be numeric 
  tmp <- data.matrix(otu_selected[, -1])
  ## 5584 * 68
  otu_selected_metabolics <- cbind(otu_selected[, 1], tmp)
  names(otu_selected_metabolics) <- names(otu_selected)
  # save(otu_selected_metabolics, file = "./RData/otu_selected_metabolics.RData")
  return(otu_selected_metabolics)
}

otu_selected_metabolics <- out_datClean()

## check zeros OTU 
cout_zeros <- function(s) {
  ## return number of zeros of vector s 
  sum(as.numeric(s) == 0)
}

## get otu as matrix 
otu_mat <- data.matrix(otu_selected_metabolics[, -1])
rownames(otu_mat) <- otu_selected_metabolics$`#NAME`
```

# Data from 2015 Mead Nitrogen Field Study

- 67 observations across treatment (high N, low N), time (July, September), and genotype
- 1--2 replicates per genotype, treatment, time combination
- 8 metabolite responses
- 5584 OTUs, some with no/few counts across all observations

```{r preproc}
## get total number of zeros for each otu 
num_zeros_per_otu <- apply(otu_mat, 1, cout_zeros)
nsamples <- ncol(otu_mat)
qplot(x = 1 - num_zeros_per_otu / nsamples, bins = 30,
     xlab = 'Proportion of non-zero samples (OTU-wise)')
qplot(x = num_zeros_per_otu / nsamples, bins = 30,
     xlab = 'Proportion of zero samples (OTU-wise)')
```

# OTU Preprocessing

- Following analyses remove OTUs with few counts or too many zeros. Exclude OTU $j$ if
    - $\le 10\%$ of observations are non-zero in OTU $j$
- Removed 1340 OTUs
- Resulting OTU matrix $\bm{X}$ has entries $[X]_{ij}$ denoting the observed count for
    - Observation $i = 1, \dots, 67$ (row)
    - OTU ID $j = 1, \dots, 4244$ (column)


```{r filter}
## get the OTU sample names 
mb_dat_sample_info$sample_name_otu <- gsub("SAMPLE_", "", mb_dat_sample_info$sample_name)
## check OTU sample names and metabolites names same 
#all(colnames(otu_mat) %in% mb_dat_sample_info$sample_name_otu)

## Create design matrix.
Trt <- mb_dat_sample_info %>% .[, 'collection_site_or_growth_conditions']
HighN <- as.numeric(Trt == 'Mead, NE - high nitrogen')
Fertilizer <- ifelse(HighN == 1, 'High Nitrogen', 'Low Nitrogen')
Date <- mb_dat_sample_info %>% .[, 'collection_month']
July <- as.numeric(Date == 'July')
Genotypes <- mb_dat_sample_info %>% .[, 'genotype']
X.otu <- t(otu_mat)
# remove columns with too many zeros
propnonzero <- 1 - num_zeros_per_otu / nsamples
keep <- (propnonzero > 0.1)
X.otu_ <- X.otu[,keep]
X <- cbind(X.otu_, HighN, July)
```

# Filtered OTU Matrix - Principal Components Analysis

First two PCs explain 35% of the variability in OTU counts.

```{r PCA}
X.otu_PCA <- prcomp(X.otu_,
                    center = TRUE,
                    scale. = TRUE)
pd <- data.frame(PropVar = summary(X.otu_PCA)$importance['Proportion of Variance',],
                 PC = 1:nsamples)
qplot(x = PC, y = PropVar, data = pd,
      xlab = 'Principal Component', ylab = 'Proportion of Variance Explained')
```

# Filtered OTU Matrix - Principal Components Analysis

First two PCs separate observations by time and treatment.

```{r PCAplot}
qplot(x = X.otu_PCA$x[,1], y = X.otu_PCA$x[,2],
      color = Fertilizer, shape = Date,
      xlab = 'PC1', ylab = 'PC2') +
    geom_point(size = 1.5) +
    scale_colour_brewer(type = 'qual', palette = 'Dark2')
```


# OTU PCA Conditional on Date, Fertilizer

```{r PCAconditional}
grp <- as.factor(Fertilizer):as.factor(Date)
cPCA <- lapply(levels(grp), function(g) {
  x <- X.otu_[grp == g,]
  Genotype <- Genotypes[grp == g]
  res <- X.otu_PCA <- prcomp(x[,-caret::nearZeroVar(x)], center = TRUE, scale. = TRUE)
  vres <- data.frame(PropVar = summary(res)$importance['Proportion of Variance',],
                     PC = 1:length(Genotype),
                     Group = g)
  out <- data.frame(PC1 = res$x[,1], 
                    PC2 = res$x[,2],
                    Group = g,
                    Genotype = Genotype)
  out$Date <- sapply(out$Group, function(x) ifelse(grepl('September', x), 'September', 'July'))
  out$Fertilizer <- sapply(out$Group, function(x) ifelse(grepl('High', x), 'High Nitrogen', 'Low Nitrogen'))
  vres$Date <- sapply(vres$Group, function(x) ifelse(grepl('September', x), 'September', 'July'))
  vres$Fertilizer <- sapply(vres$Group, function(x) ifelse(grepl('High', x), 'High Nitrogen', 'Low Nitrogen'))
 return(list(out, vres))
})


cPCApd <- do.call('rbind', lapply(cPCA, function(res) res[[1]]))
cPCApd$Breeding <- ifelse(cPCApd$Genotype %in% c('CU-C126', 'CU-C225', 'CU-CO53', 'CU-CO56'), 'Hybrid', 'Inbred')
cPCApdSept <- cPCApd %>% filter(Date == 'September')
cPCAve <- do.call('rbind', lapply(cPCA, function(res) res[[2]]))
cPCAveSept <- cPCAve %>% filter(Date == 'September')
qplot(x = PC, y = PropVar, data = cPCAve,
      xlab = 'Principal Component', ylab = 'Proportion of Variance Explained') +
  facet_wrap(~Group, ncol = 2)
qplot(x = PC, y = PropVar, data = cPCAveSept,
      xlab = 'Principal Component', ylab = 'Proportion of Variance Explained') +
  facet_wrap(~Fertilizer, ncol = 2)
```

# OTU PCA Conditional on Date, Fertilizer

```{r PCAcondplot}
qplot(x = PC1, y = PC2, color = Genotype, shape = Breeding, data = cPCApd) +
    geom_point(size = 2.5) +
    facet_wrap(~Group, ncol = 2) 

qplot(x = PC1, y = PC2, color = Genotype, shape = Breeding, data = cPCApdSept) +
    geom_point(size = 2.5) +
    facet_wrap(~Fertilizer, ncol = 2) 
```

# OTU Pairwise Correlations

```{r corr}
# pairwise correlations -- function to plot
corploter <- function(A, Date, Fertilizer, approx.nviz = 200, method = 'spearman') {
    timelevels <- unique(Date)
    trtlevels <- unique(Fertilizer)
    colnames(A) <- sapply(colnames(A), function(x) gsub('[[:alpha:]]+_', '', x))
    R <- cor(A, method = method)
    dR <- reshape2::melt(R); dR$cond <- 'Pooled Nitrogen'; dR$Date <- 'Pooled Date'
    dRupper.tri <- dR %>% filter(Var1 < Var2) %>% mutate(rank = min_rank(desc(abs(value))))
    dRupper.trikeep <- dRupper.tri %>% filter(rank < approx.nviz*1.28)
    keepOTU <- unique(c(dRupper.trikeep$Var1, dRupper.trikeep$Var2))
    message(paste('Plotting', length(keepOTU), 'top marginally correlated OTUs'))
    dR <- dR %>% filter(Var1 %in% keepOTU & Var2 %in% keepOTU)
    R1 <- cor(A[Fertilizer == trtlevels[1], colnames(A) %in% keepOTU], method = method)
    dR1 <- reshape2::melt(R1); dR1$cond <- trtlevels[1]; dR1$Date <- 'Pooled Date'
    R0 <- cor(A[Fertilizer == trtlevels[2], colnames(A) %in% keepOTU], method = method)
    dR0 <- reshape2::melt(R0); dR0$cond <- trtlevels[2]; dR0$Date <- 'Pooled Date'
    Rcond <- lapply(timelevels, function(thisdate) {
        D <- Date == thisdate
        Rd <- cor(A[D, colnames(A) %in% keepOTU], method = method)
        dRd <- reshape2::melt(Rd); dRd$cond <- 'Pooled Nitrogen'
        Rd1 <- cor(A[Fertilizer == trtlevels[1] & D, colnames(A) %in% keepOTU], method = method)
        dRd1 <- reshape2::melt(Rd1); dRd1$cond <- trtlevels[1]
        Rd0 <- cor(A[Fertilizer == trtlevels[2] & D, colnames(A) %in% keepOTU], method = method)
        dRd0 <- reshape2::melt(Rd0); dRd0$cond <- trtlevels[2]
        out <- rbind(dRd, dRd1, dRd0)
        out$Date <- thisdate
        return(out)
    })
    dRcond <- do.call('rbind', Rcond)
    cordat <- rbind(dRcond, dR, dR0, dR1)
    cordat$cond <- relevel(factor(cordat$cond), ref = 'Pooled Nitrogen')
    cordat$Date <- relevel(factor(cordat$Date), ref = 'Pooled Date')
    cordat <- cordat %>% mutate(Var1 = as.factor(Var1), Var2 = as.factor(Var2))
    return(cordat)
}

o <- corploter(X.otu_, Date, Fertilizer)
ggplot(o) +
    geom_tile(aes(x = Var1, y = Var2, fill = value), color = 'white') +
    facet_grid(Date~cond) + 
    scale_y_discrete(limits = rev(levels(o$Var2))) +
    scale_fill_gradient2(low = "orange", high = "purple", mid = "white",  
                         midpoint = 0, limit = c(-1,1), space = "Lab", 
                         name="Spearman\nCorrelation") +
    xlab('') + ylab('') +
    theme(axis.text.x = element_blank(), axis.text.y = element_blank())
# september only
oSept <- o %>% filter(Date == 'September')
as.nf <- function(x) as.numeric(levels(x))[x]
keepOTU <- unique(oSept$Var1[!is.na(oSept$value) & oSept$cond == 'Pooled Nitrogen' & as.nf(oSept$Var1) > as.nf(oSept$Var2)])
oSept <- oSept %>% filter(Var1 %in% keepOTU & Var2 %in% keepOTU)
ggplot(oSept) +
    geom_tile(aes(x = Var1, y = Var2, fill = value), color = 'white') +
    facet_grid(~cond) + 
    scale_y_discrete(limits = rev(levels(oSept$Var2))) +
    scale_fill_gradient2(low = "orange", high = "purple", mid = "white",  
                         midpoint = 0, limit = c(-1,1), space = "Lab", 
                         name="Spearman\nCorrelation") +
    xlab('') + ylab('') +
    theme(axis.text.x = element_blank(), 
          axis.text.y = element_blank(),
          legend.position = 'bottom',
          legend.key.size = unit(1, 'cm'))
ggsave('corr-sept.pdf', width = 10, height = 5, dpi = 320)
ggsave('corr-sept.png', width = 10, height = 5, dpi = 320)
# July only
oJuly <- o %>% filter(Date == 'July')
as.nf <- function(x) as.numeric(levels(x))[x]
keepOTU <- unique(oJuly$Var1[!is.na(oJuly$value) & oJuly$cond == 'Pooled Nitrogen' & as.nf(oJuly$Var1) > as.nf(oJuly$Var2)])
oJuly <- oJuly %>% filter(Var1 %in% keepOTU & Var2 %in% keepOTU)
ggplot(oJuly) +
    geom_tile(aes(x = Var1, y = Var2, fill = value), color = 'white') +
    facet_grid(~cond) + 
    scale_y_discrete(limits = rev(levels(oSept$Var2))) +
    scale_fill_gradient2(low = "orange", high = "purple", mid = "white",  
                         midpoint = 0, limit = c(-1,1), space = "Lab", 
                         name="Spearman\nCorrelation") +
    xlab('') + ylab('') +
    theme(axis.text.x = element_blank(), 
          axis.text.y = element_blank(),
          legend.position = 'bottom',
          legend.key.size = unit(1, 'cm'))
ggsave('corr-july.pdf', width = 10, height = 5, dpi = 320)
ggsave('corr-july.png', width = 10, height = 5, dpi = 320)
```


# Analysis for September

```{r sept}
require(SISmod)
require(hdi)
otu_names <- colnames(X.otu_)
p <- length(otu_names)
A1 <- SISmod::scale(X.otu_[Fertilizer == 'High Nitrogen' & Date == 'September',])
A0 <- SISmod::scale(X.otu_[Fertilizer == 'Low Nitrogen' & Date == 'September',])
colnames(A1) <- colnames(A0) <- 1:p
# check samples are in same order in y and X
all(rownames(X.otu_) == sapply(mb_dat_sample_info$sample_name, function(x) gsub('SAMPLE_', '', x)))

# responses -- look at each metabolite
## Merge metabolite data with sample info, sort to match OTU order
mb_dat_merge <- left_join(mb_dat, mb_dat_sample_info, by = c("Sample" = "sample_name_metabolites")) %>% data.frame()
ord <- match(rownames(X.otu_), mb_dat_merge$sample_name_otu)
## Grab responses, log transform, and put them in a list
Ys <- lapply(names(mb_dat)[-1], function(nm) {
  idx <- mb_dat_sample_info %>% .[, 'sample_name_metabolites']
  out <- mb_dat %>% filter(Sample %in% idx) 
  y <- data.frame(y = out[ord,nm], ID = out[ord, 'Sample'])
  names(y) <- c('y', 'ID')
  return(y)
})
names(Ys) <- names(mb_dat)[-1]


fit <- function(x, y, ID, tune = 'bic', std = FALSE) {
    yc <- (y - mean(y))
    n <- length(y); p <- ncol(x)
    ISISlasso <- tryCatch(suppressWarnings(SIS(x = x, y = y, penalty = 'lasso', 
                                               tune = tune, nfolds = n, , standardize = std)),
                          error = function(e) NA)
    b.ISISlasso <- ISISlasso$coef.est[-1]
    idx <- as.numeric(sapply(names(b.ISISlasso), function(s) gsub('X', '', s)))
    x.ISISlasso <- x[,idx]
    if (class(x.ISISlasso) != 'matrix') x.ISISlasso <- matrix(x.ISISlasso, ncol = 1)
    sigma.ISISlasso <- sqrt(sum((yc - x.ISISlasso%*%b.ISISlasso)^2) / (n - length(b.ISISlasso) - 1)) # naive est of sigma
    ISISlasso.bh <- tryCatch(suppressWarnings(lasso.proj(x = x.ISISlasso, y = y, standardize = std, 
                                                         suppress.grouptesting = TRUE, sigma = sigma.ISISlasso,
                                                         multiplecorr.method = 'BH', betainit = b.ISISlasso, return.Z = TRUE)), error = function(e) NA)
    if (!is.na(ISISlasso.bh[[1]])) {
        yhat <- as.numeric(x.ISISlasso %*% ISISlasso.bh$bhat + mean(y))
    } else {
        yhat <- rep(mean(y), n)
    }
    ISISlasso.bh$resid <- y - yhat
    ISISlasso.bh$yhat <- yhat
    ISISlasso.bh$SampleID <- ID
    return(ISISlasso.bh)
}

# extract results
sumdatcond <- function(res0, res1, p, alpha = 1) {
    b1 <- b0 <- rep(0, p)
    betahat1 <- betahat0 <- rep(0, p)
    se1 <- se0 <- rep(0, p)
    if (class(res0)[1] == 'logical') {
        b0 <- betahat0 <- rep(NA, p)
    } else {
        ids0 <- as.numeric(names(res0$pval))
        b0[ids0] <- res0$bhat
        betahat0[ids0] <- res0$betahat
        se0[ids0] <- res0$se
    }
    if (class(res1)[1] == 'logical') {
        b1 <- betahat1 <- rep(NA, p)
    } else {
        ids1 <- as.numeric(names(res1$pval))
        b1[ids1] <- res1$bhat
        betahat1[ids1] <- res1$betahat
        se1[ids1] <- res1$se
    }
    b <- (b0 + b1)
    betahat <- (betahat0 + betahat1 ) / 2
    se <- sqrt(se0^2 + se1^2)
    z <- b / se
    ps <- 2 * (1 - pnorm(abs(z)))
    ps[is.na(ps)] <- 1
    ps1 <- 2 * (1 - pnorm(abs(b1 / se1)))
    ps1[is.na(ps1)] <- 1
    ps0 <- 2 * (1 - pnorm(abs(b0 / se0)))
    ps0[is.na(ps0)] <- 1
    pscorr <- p.adjust(ps, method = 'fdr')
    pscorr0 <- p.adjust(ps0, method = 'fdr')
    pscorr1 <- p.adjust(ps1, method = 'fdr')
    df <- data.frame(OTU = rep(otu_names, times = 3), 
                     betahat = c(betahat, betahat0, betahat1), 
                     pval = c(ps, ps0, ps1), 
                     pval.corr = c(pscorr, pscorr0, pscorr1),
                     condition = rep(c('Standardized', 'Low Nitrogen', 'High Nitrogen'), each = p))
    if (alpha < 1) {
        keepOTUs <- df$OTU[df$pval.corr <= alpha & df$condition == 'Standardized']
        df <- df %>% filter(OTU %in% keepOTUs)
    }
    names(df) <- c('Feature', 'Estimate', 'p-value', 'BH q-value', 'Fertilizer')
    return(df)
}
```

# Results (analysis on original scale)

```{r res, message = FALSE, warming = FALSE}
fit1 <- lapply(Ys, function(y) fit(A1, 
                                   y$y[Fertilizer == 'High Nitrogen' & Date == 'September'], 
                                   y$ID[Fertilizer == 'High Nitrogen' & Date == 'September']))
fit0 <- lapply(Ys, function(y) fit(A0, 
                                   y$y[Fertilizer == 'Low Nitrogen' & Date == 'September'],
                                   y$ID[Fertilizer == 'Low Nitrogen' & Date == 'September']))
# check that variables were selected in at least one condition
sapply(fit1, function(x) class(x) == 'logical') + sapply(fit0, function(x) class(x) == 'logical')
ans <- lapply(1:length(Ys), function(j) sumdatcond(fit0[[j]], fit1[[j]], p, alpha = 0.05))
labs <- sapply(names(Ys), function(x) str_to_title(gsub('_', ' ', x)))
labs[labs == '12 Oxo Phytodienoic Acid'] <- '12-oxo Phytodienoic Acid'
names(ans) <- labs
resids <- lapply(1:length(Ys), function(j) {
    f0 <- fit0[[j]]; f1 <- fit1[[j]]
    pd <- data.frame(e = c(f0$resid, f1$resid),
                     yhat = c(f0$yhat, f1$yhat),
                     SampleID = c(f0$SampleID, f1$SampleID),
                     condition = c(rep('Low Nitrogen', length(f0$resid)), rep('High Nitrogen', length(f1$resid))))
    pd$Metabolite <- labs[j]
    out <- ggplot(pd, aes(x = yhat, y = e, color = condition)) + 
        geom_point() +
        xlab('Conditional Predicted Value') + ylab('Residual') + labs(color = 'Fertilizer') +
        ggtitle(labs[j]) +
        scale_colour_brewer(type = 'qual', palette = 'Dark2')
    qq <- ggplot(pd, aes(sample = e)) +
        stat_qq() + stat_qq_line() +
        facet_grid(~condition) +
        xlab('Theoretical') + ylab('Sample') +
        ggtitle(labs[j])
    ans <- list(residplt = out, qqplot = qq, pd = pd)
    return(ans)
})

```

```{r printres, results = 'asis'}
require(pander)
panderOptions('knitr.auto.asis', FALSE)
for (j in 1:length(Ys)) {
    cat('\n')
    cat(paste('##', labs[j], '\n'))
    pander(ans[[j]])
    cat('\n')
    print(resids[[j]]$residplt)
    print(resids[[j]]$qqplot)
    cat('\n')
}

residdat <- do.call('rbind', lapply(resids, function(r) r$pd))
# sample name of greatest residual
maxresids <- residdat %>% 
    group_by(Metabolite) %>% 
    summarise(MaxAbsResid = SampleID[which.max(abs(e))])
pander(maxresids)


# output tables
dir.create('./2015-DOE-metabolite-microbiome_tables')
for (i in 1:length(ans)) {
    tbl <- xtable::xtable(ans[[i]], display = c('g', 's', 'g', 'g', 'g', 's'),  
                          digits = 2, math.style.exponents = TRUE)
    print(tbl, file = paste0('./2015-DOE-metabolite-microbiome_tables/', labs[i], '.tex'))
}
```

```{r corODPA}
OTUS <- ans[[1]] %>% filter(`BH q-value` < 0.05 & Fertilizer == 'Standardized')
A <- cbind(X.otu_[, colnames(X.otu_) %in% OTUS$Feature], Ys[[1]]$y)
colnames(A) <- c(as.character(OTUS$Feature), 'OPDA')
trtlevels <- unique(Fertilizer)
R <- cor(A[Date == 'September', ], method = 'spearman')
dR <- reshape2::melt(R); dR$cond <- 'Pooled Nitrogen'
R1 <- cor(A[Date == 'September' & Fertilizer == trtlevels[1], ], method = 'spearman')
dR1 <- reshape2::melt(R1); dR1$cond <- trtlevels[1]
R0 <- cor(A[Date == 'September' & Fertilizer == trtlevels[2], ], method = 'spearman')
dR0 <- reshape2::melt(R0); dR0$cond <- trtlevels[2]
cordat <- rbind(dR, dR0, dR1)
cordat$cond <- relevel(factor(cordat$cond), ref = 'Pooled Nitrogen')
ggplot(cordat) +
    geom_tile(aes(x = Var1, y = Var2, fill = value), color = 'white') +
    facet_grid(~cond) + 
    scale_y_discrete(limits = rev(levels(cordat$Var2))) +
    scale_fill_gradient2(low = "orange", high = "purple", mid = "white",  
                         midpoint = 0, limit = c(-1,1), space = "Lab", 
                         name="Spearman\nCorrelation") +
    xlab('') + ylab('') +
    theme(legend.position = 'bottom',
          legend.key.size = unit(1, 'cm'))

# summary stats for low vs high N for selected OTUs
dfA <- data.frame(A, Date, Fertilizer)
#min
dfA %>% 
    group_by(Date, Fertilizer) %>%
    summarize(OTU_115 = min(OTU_115),
              OTU_385 = min(OTU_385),
              OTU_409 = min(OTU_409),
              OTU_855 = min(OTU_855)) %>% 
    kable()
#max
dfA %>% 
    group_by(Date, Fertilizer) %>%
    summarize(OTU_115 = max(OTU_115),
              OTU_385 = max(OTU_385),
              OTU_409 = max(OTU_409),
              OTU_855 = max(OTU_855)) %>%
    kable()
# mean (stdev)
 meanSd <- function(x, digits = 2) {
     xmean <- mean(x)
     xsd <- sd(x)
     return(paste0( round(xmean, digits), ' (', round(xsd, digits), ')', collapste = ''))
 }
 dfA %>% 
    group_by(Date, Fertilizer) %>%
    summarize(OTU_115 = meanSd(OTU_115),
              OTU_385 = meanSd(OTU_385),
              OTU_409 = meanSd(OTU_409),
              OTU_855 = meanSd(OTU_855)) %>%
     kable()
```
````
